# An Unconventional Targeting Architecture Directs Cardiolipin Synthase to Multiple Compartments

**DOI:** 10.64898/2026.11.24.768828

**Authors:** Swapnil Sahoo, Jan Janouškovec, Abdur Rahaman

## Abstract

Multi-compartment protein targeting underlies the functional and metabolic flexibility of eukaryotic cells. However, how a single targeting region directs a protein to multiple cellular compartments remains poorly understood. A good example is membrane lipid biosynthesis: cardiolipin is normally synthesized in the mitochondrial inner membrane but has also been detected in the nuclear envelope of *Tetrahymena*. Here, we show that *Tetrahymena* cardiolipin synthase (TtCLS) localises to three distinct compartments. Systematic evolutionary and cell-biological analyses uncover a lineage-specific C-terminal targeting signal that expands the subcellular distribution of TtCLS across mitochondria, nuclear envelope and plasma membrane. We further show that its dynamic distribution among these compartments is regulated by phosphorylation. This multifunctional targeting architecture contains an amphipathic helix implicated in membrane association, which is structurally conserved across bacterial and eukaryotic CLS_PLD enzymes. Notably, this architecture remains compatible with conserved mitochondrial import systems across evolutionarily distant eukaryotes. Together, these findings uncover a previously unknown targeting architecture in which a single C-terminal region integrates overlapping and potentially regulatable localization determinants to target one protein across three different membrane systems.

## Introduction

Protein-targeting signals are generally viewed as discrete sequence elements that direct proteins to defined intracellular destinations. Nuclear-encoded proteins are synthesized in the cytosol and subsequently sorted to their respective cellular compartments through specific targeting signals recognized by dedicated trafficking and translocation machineries^1^. Although individual proteins are often thought to have a fixed cellular localization, growing evidence indicates that their targeting can be regulated, allowing them to access different cellular compartments^1–3^. Such differential targeting expands the functional repertoire of proteins without increasing genome complexity and provides a means of coordinating distinct cellular processes through regulated spatial distribution^3,4^.

Among intracellular trafficking pathways, mitochondrial protein import is one of the best understood. Most nuclear-encoded mitochondrial proteins possess N-terminal amphipathic mitochondrial targeting sequences (MTSs) that mediate recognition by the conserved TOM and TIM translocase complexes^5–8^. Nevertheless, mitochondrial targeting is considerably more diverse than once appreciated^7–9^. Members of the mitochondrial carrier family rely on multiple internal targeting elements distributed throughout the protein rather than on a cleavable N-terminal pre-sequence^10^, whereas tail-anchored proteins such as hFis1 employ C-terminal targeting information for localization to both the mitochondrial outer membrane and the peroxisomal membrane^11,12^. Likewise, several proteins partition between mitochondria and other organelles through overlapping or competing targeting signals whose activities are regulated by mechanisms such as phosphorylation^3^.

Despite these advances^3,4^, the diversity of signals specifying multiple cellular destinations remains poorly explored. For C-terminal regions, experimentally characterized examples of naturally occurring dual organelle targeting identified to date come from animals, plants and fungi, and involve integral membrane proteins^12–14^. How a single targeting region can distribute proteins among three or more cellular compartments remains poorly resolved, despite the importance of flexible enzyme localization for cellular metabolism. This is particularly relevant to lipid metabolic enzymes, including cardiolipin synthases, because localized lipid synthesis directly contributes to membrane identity and organelle function^15–18^.

Ciliates are unusual among eukaryotes in retaining a bacterial-type phospholipase D cardiolipin synthase (CLS_PLD), whereas most eukaryotes use the distinct CDP-alcohol phosphatidyltransferase-type cardiolipin synthase (CLS_CAP)^19^. Cardiolipin is the signature phospholipid of mitochondria, where it is essential for maintaining cristae architecture, respiratory chain organization, membrane dynamics and oxidative phosphorylation^20–22^. However, cardiolipin has also been detected at the nuclear envelope of *Tetrahymena*^23^, suggesting that cardiolipin metabolism extends beyond mitochondria. In fact, cardiolipin has been shown to be important for macronuclear expansion during sexual reproduction in *Tetrahymena thermophila*^24^. Similarly, extra-mitochondrial cardiolipin and peripheral localization of CLS_PLD have been reported during specific developmental stages of *Plasmodium*^25^. These observations raise the possibility that CLS_PLD proteins can access membrane compartments beyond mitochondria. However, whether such extra-mitochondrial distribution is developmentally regulated, and how this targeting information is encoded in the sequence, remain unknown.

Here, we demonstrate that *Tetrahymena* cardiolipin synthase employs an unconventional targeting mechanism centred on a C-terminal localization signal. Rather than functioning solely as a mitochondrial targeting sequence, this signal region contains overlapping localization determinants that direct TtCLS to the mitochondrial inner membrane, nuclear envelope and plasma membrane. Systematic truncation analysis revealed that these determinants are integrated within a compact C-terminal region and that their localization outputs are shaped by the surrounding sequence context rather than functioning as independent targeting motifs. Site-directed mutagenesis further implicated phosphorylation as a mechanism for modulating the distribution of TtCLS among these compartments. Despite lacking a canonical N-terminal mitochondrial targeting sequence, TtCLS appears to be sorted to the inner mitochondrial membrane via the matrix, and is targeted to mitochondria in mammalian cells. Together, our findings reveal unexpected plasticity in protein-targeting signal architecture and identify a multifunctional C-terminal localization signal as a previously unrecognized strategy for coordinating protein localization across multiple membrane compartments while remaining compatible with conserved intracellular trafficking machinery.

## Results

### 1. A single cardiolipin synthase localizes to three distinct compartments in *Tetrahymena* despite lacking canonical N-terminal targeting sequences

Sequence similarity searches against the *Tetrahymena thermophila* SB210 genome identified a single CLS_PLD gene (*TtCLS*) and no CLS_CAP cardiolipin synthase genes. To determine the intracellular localization of TtCLS, we C-terminally tagged its endogenous locus with 3×mCherry by homologous recombination, preserving its native genomic promoter.

Confocal microscopy revealed that endogenously expressed TtCLS localized predominantly to mitochondria (**Fig. 1**), consistent with its established role in cardiolipin biosynthesis. Surprisingly, in addition to the mitochondrial signal, TtCLS was also detected at the nuclear envelope and plasma membrane (**Fig. 1**). The nuclear envelope localization is consistent with previous reports describing cardiolipin at this membrane^23^. Together, these observations demonstrate that the only cardiolipin synthase of *Tetrahymena* is targeted to three distinct membrane compartments. This unexpected localization pattern prompted us to investigate the molecular mechanism underlying the compartment-specific targeting of TtCLS.

**Figure 1:**
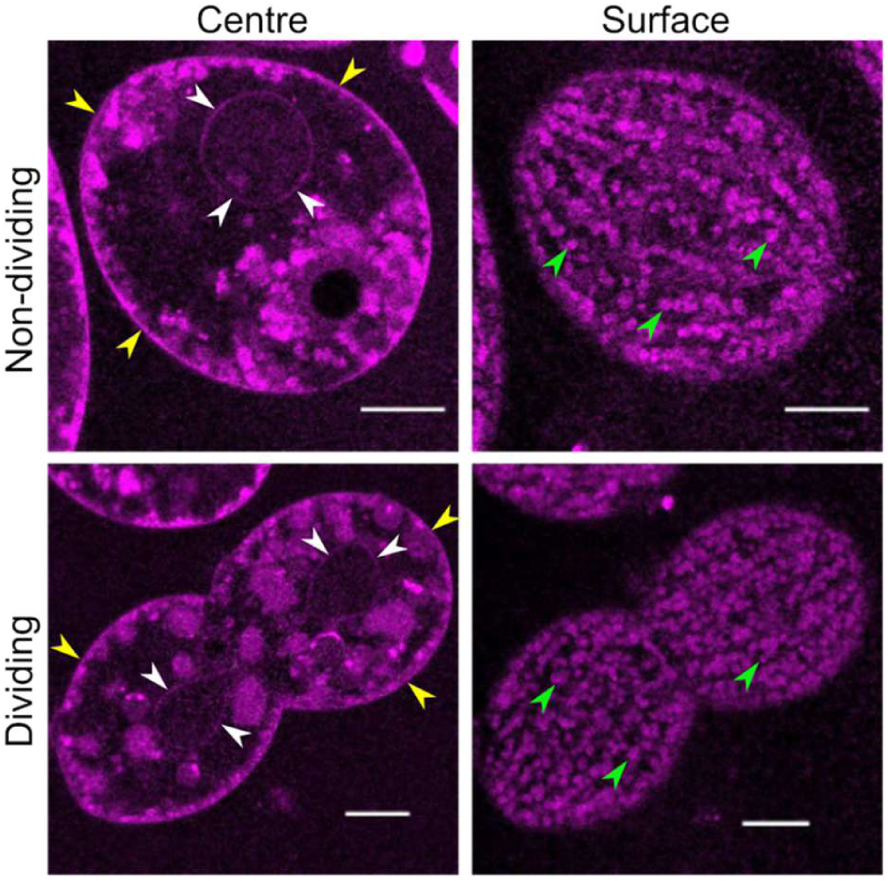
*Tetrahymena* cardiolipin synthase localizes to mitochondria, the nuclear envelope and the plasma membrane. Representative Confocal images of a live *Tetrahymena* cell expressing endogenously tagged cardiolipin synthase from n= 3 independent experiments. While white arrowhead indicates nuclear envelope, yellow arrowhead shows plasma-membrane localisation (left panel). Right panel depicts cortical region showing mitochondrial localization indicated by green arrowhead. Scale bar-10 μm.

Most nuclear-encoded mitochondrial proteins are targeted through an N-terminal mitochondrial targeting sequence (MTS) recognized by the mitochondrial protein import machinery^7,26^. We therefore examined whether TtCLS contains canonical sequence features that could explain its localization to mitochondria, the nuclear envelope or plasma membrane. Its amino acid sequence was analysed using multiple prediction algorithms, including TargetP 2.0^27^, SignalP 6.0^28^, DeepLoc 2.0^29^, DeepMito^30^ and WoLF PSORT^31^ (**Table S1**). None of the prediction tools identified recognizable localization signals that could account for the experimentally observed localization of TtCLS to mitochondria, the nuclear envelope or the plasma membrane. These observations suggest that the intracellular distribution of TtCLS is governed by an unconventional targeting mechanism that is not detected by existing prediction algorithms.

### 2. The mitochondrial targeting information resides within the C-terminal half of TtCLS

To identify the sequence responsible for mitochondrial targeting, the full-length TtCLS was expressed as a fusion protein with GFP at the N-terminus (GFP-TtCLS) and its intracellular localization was examined by confocal microscopy. Similar to the endogenously tagged protein, overexpressed GFP-TtCLS localized predominantly to mitochondria (**Fig. 2A**). In contrast to the endogenously-tagged protein, however, we were unable to detect localization at either the nuclear envelope or the plasma membrane. This difference may reflect altered accessibility or regulation of the localization signals under overexpression conditions or difference in tag type and position. Nevertheless, the exclusive mitochondrial localization of the overexpressed protein provided a suitable system for systematically mapping the mitochondrial targeting sequence.

**Figure 2:**
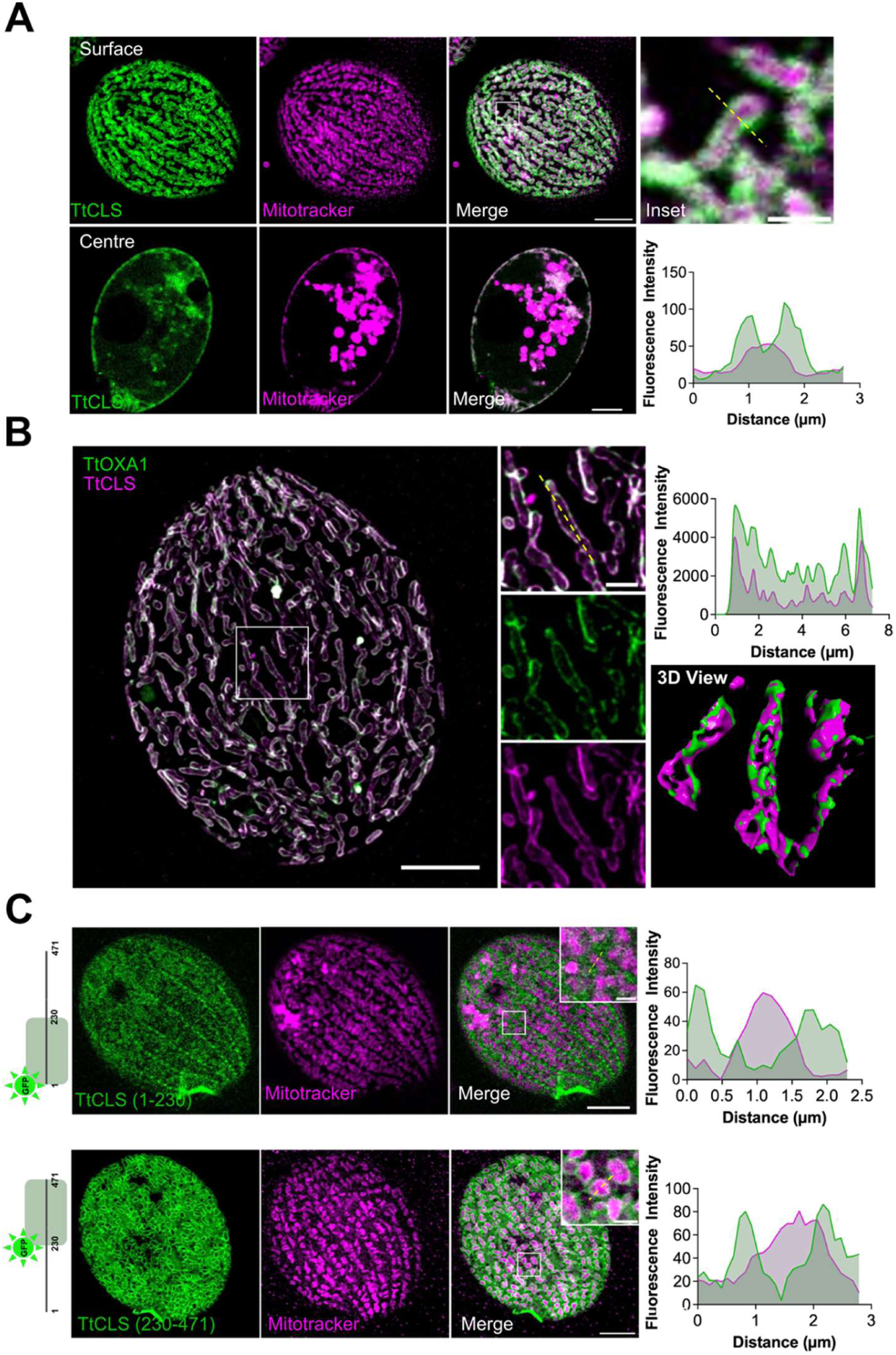
A C-terminal region directs TtCLS to the mitochondrial inner membrane. **(A)** Confocal images of live *Tetrahymena* cells expressing either expressing GFP–TtCLS after incubation with mitotracker deep red. **(B)** SR-SIM² image of a *Tetrahymena* cell co-expressing GFP–OXA1 and mCherry–TtCLS with a 3D view. **(C)** Confocal images of *Tetrahymena* cells expressing the fragment encompassing 1-230 amino acid residues (TtCLS 1-230) or 231–471 amino acid residues as N-terminally tagged GFP fusion protein after incubation with mitotracker deep red. The boxed region is enlarged in the inset. Corresponding fluorescence intensity profiles and the schematic of the fragments are shown in the right. Scale bars, 1 μm (inset) and10 μm (main images). Data are representative of n= 3 independent experiments, >30 cells per experiment, and >20 mitochondria per cell.

To determine the sub-mitochondrial localization of TtCLS, we first characterized the localization of TtOXA1, for use as an inner mitochondrial membrane marker. This conserved eukaryotic insertase was previously identified in the *Tetrahymena* mitochondrial proteome^32^. N-terminally GFP-tagged TtOXA1 was expressed and its localization was assessed relative to MitoTracker-labelled mitochondria. Confocal microscopy revealed that GFP-TtOXA1 localized to mitochondria and had a membrane-associated distribution consistent with its expected localization to the inner mitochondrial membrane (**Fig. S1A**). Super-resolution Structured Illumination Microscopy (SR-SIM^2^) demonstrated extensive colocalization between mCherry-TtCLS and GFP-TtOXA1 throughout the mitochondrial network (**Fig. 2B**). At higher magnification, both proteins decorated the internal mitochondrial membrane architecture including cristae, consistent with localization to the inner mitochondrial membrane (**Fig. 2B**). To further evaluate this localization, fluorescence intensity profiles were generated along individual mitochondria. The fluorescence intensity profiles of mCherry-TtCLS and GFP-TtOXA1 exhibited closely matching peak distributions across the length of the mitochondria, with multiple coincident local maxima corresponding to internal membrane structures resolved by SR-SIM^2^ (**Fig. 2B**). The extensive overlap between the two fluorescence profiles is consistent with enrichment of TtCLS along the inner mitochondrial membrane, in agreement with its expected role in cardiolipin biosynthesis. Moreover, the 3D reconstruction of the SR-SIM^2^ images also revealed the localisation of the TtCLS onto the cristae labelled by TtOXA1 **(Fig. 2B)**.

Having established that TtCLS localizes to the inner mitochondrial membrane, we next sought to identify the sequence responsible for mitochondrial targeting. For this purpose, the protein was divided into N-terminal and C-terminal halves, each expressed as a fusion protein with GFP at the N-terminus (**Fig. 2C**). Confocal microscopy revealed that the N-terminal half failed to localize to mitochondria and instead displayed diffuse cytoplasmic fluorescence. In contrast, the C-terminal half efficiently localized to mitochondria (**Fig. 2C**), demonstrating that the mitochondrial targeting information resides within the C-terminal region of TtCLS. These observations are consistent with our sequence analysis, which failed to identify a canonical N-terminal mitochondrial targeting sequence, and instead suggest that *Tetrahymena* cardiolipin synthase utilizes a previously unrecognized C-terminal targeting mechanism.

### 3. A C-terminal region directs TtCLS to the mitochondria

We next sought to narrow down the sequence responsible for the mitochondrial localization of TtCLS. Progressive truncation of the C-terminal region generated a fragment encompassing residues 356–471, which was expressed as a fusion protein with GFP at the N-terminus. Confocal microscopy revealed that this fragment retained robust mitochondrial localization comparable to that of the parental C-terminal construct (**Fig. 3A**), indicating that the mitochondrial targeting information is contained within the C-terminal 116 amino acids of TtCLS. Analysis of alignment conservation revealed these 116 residues still contain a part of conserved core around the second catalytic domain (as described in section 7).

**Figure 3:**
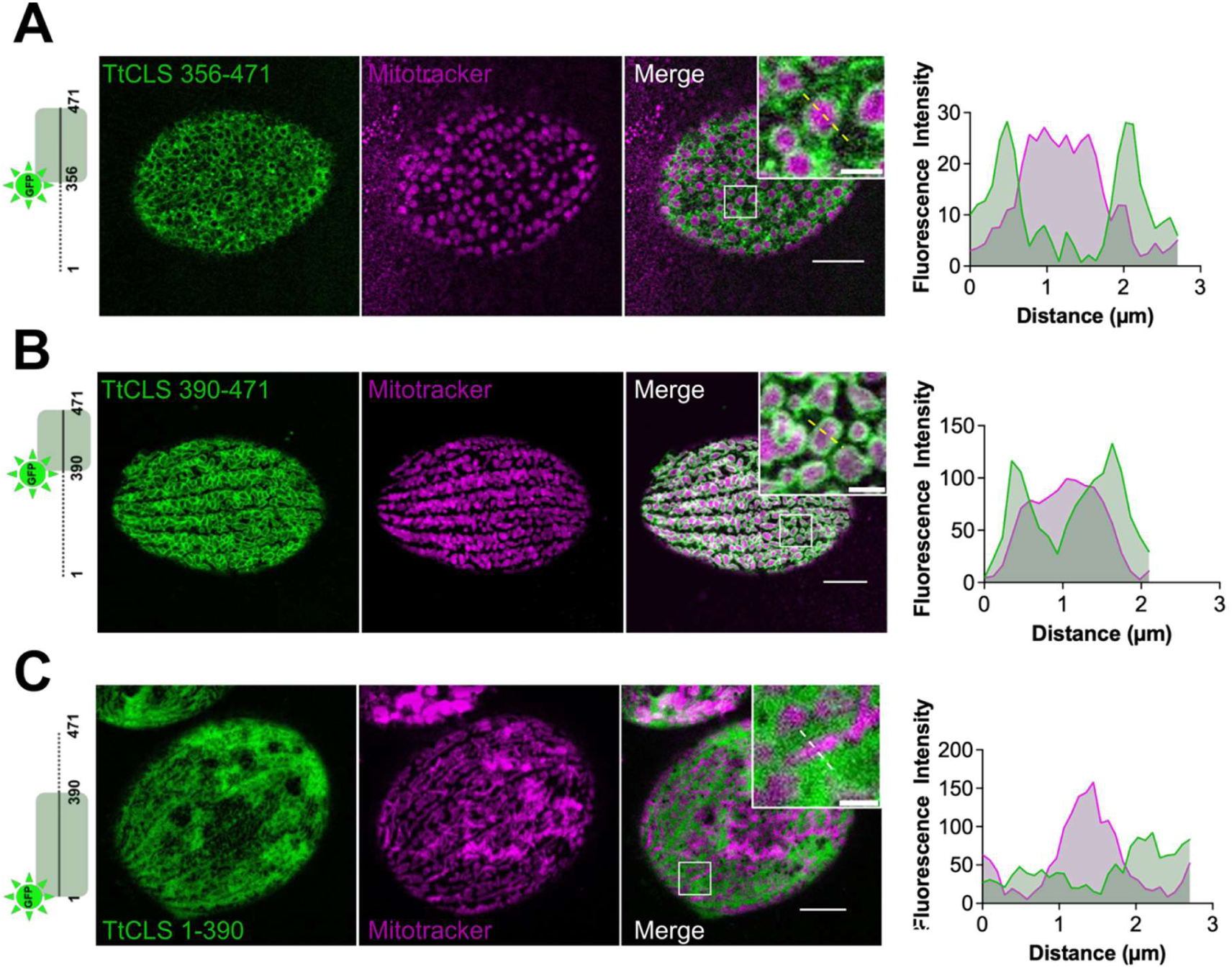
The C-terminal 82 amino acids of TtCLS are sufficient for mitochondrial targeting. **(A)** Representative Confocal images of live *Tetrahymena* cells from n= 4 independent experiments, >30 cells per experiment, and >20 mitochondria per cell, expressing GFP–TtCLS 356–471 **(B)** GFP–TtCLS 390–471 and **(C)** GFP–TtCLS 1–390 after labelling with MitoTracker deep red. Insets show respective enlarged boxed regions. Fluorescence intensity profiles are also shown. Scale bars, 1 μm (inset) and10 μm (main images).

We therefore truncated the region further and expressed the 82-amino-acid C-terminal region (390–471) as a GFP-TtCLS 390-471 fusion. Remarkably, this fragment alone was sufficient to efficiently target GFP to mitochondria, producing a localization pattern comparable to that observed for the full-length protein (**Fig. 3B**). In contrast, the fragment encompassing residues 1–390, corresponding to the conserved catalytic core of TtCLS, exhibited diffuse cytoplasmic localization with no detectable mitochondrial signal (**Fig. 3C**). Together, these findings identify the lineage-specific C-terminal sequence as the mitochondrial targeting region and is sufficient to direct mitochondrial localization independently of the conserved catalytic core.

### 4. The C-terminal localization signal contains overlapping targeting determinants for mitochondria, the plasma membrane and the nuclear envelope

Having identified the C-terminal region as sufficient for mitochondrial localization, we next sought to define the minimal sequence required for mitochondrial targeting. To this end, a series of progressive truncations was generated from both the N- and C-termini of the 390–471 amino acid localization signal, and their intracellular localization was examined by confocal microscopy.

To further refine the mitochondrial targeting sequence, we deleted N-terminal 21 amino acids (residues 390–410) from C-terminal region and expressed as GFP fusion. Surprisingly, instead of simply abolishing mitochondrial localization, the resulting fragment (411–471) was redirected exclusively to the plasma membrane and nuclear envelope (**Fig. 4A, 4B, 4C**). This unexpected observation revealed that the C-terminal localization signal contains additional targeting information that is normally masked in the presence of residues 390–410.

**Figure 4:**
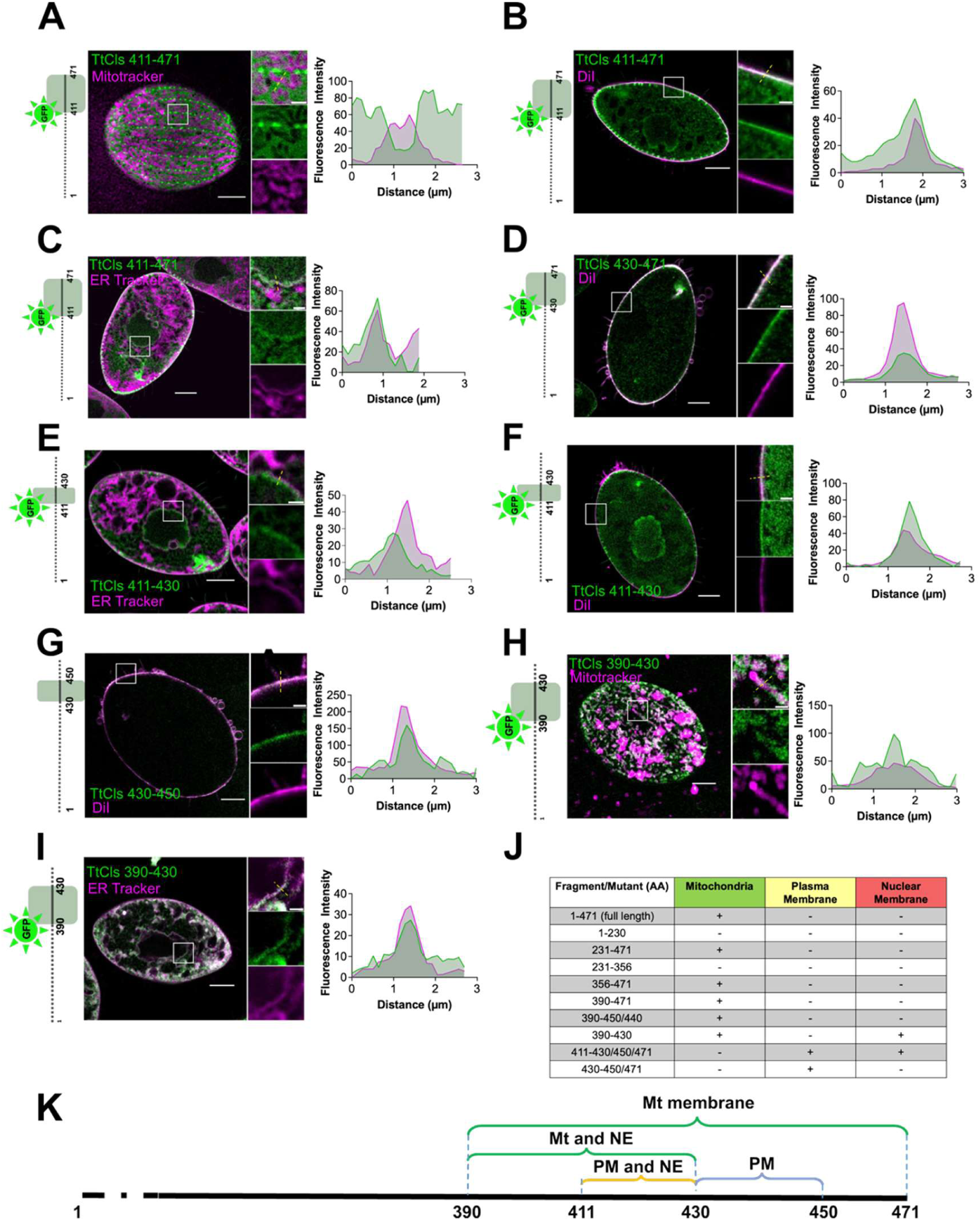
The C-terminal signal integrates overlapping targeting determinants for three membrane compartments. **(A-I)** Confocal images of *Tetrahymena* cells from n= 3 independent experiments, >30 cells per experiment, >20 mitchondria per cell, >5 regions of plasma membrane or nuclear envelope, expressing the indicated TtCLS C-terminal truncation constructs and labelled with MitoTracker (mitochondria), ER Tracker (endoplasmic reticulum) or Dil (plasma membrane). The corresponding fluorescence intensity profiles show the spatial distribution of TtCLS with respect to the indicated compartment marker. Insets show enlarged regions of the boxed areas. Main image scale bars, 10 μm; inset scale bars, 1 μm. **(J)** Schematic summary of the localization determinants identified within the C-terminal region of TtCLS. **(K)** Summary of the localization phenotypes of the indicated TtCLS fragments.

Therefore, we systematically dissected the localization signal to identify the determinants responsible for plasma membrane and nuclear envelope targeting. Removal of an additional 19 amino acids generated the 430–471 fragment, which retained plasma membrane localization but completely lost localization to the nuclear envelope (**Fig. 4D**), indicating that residues 411–429 are required for nuclear envelope targeting. Consistent with this interpretation, residues 411–430 alone were sufficient to direct GFP to the nuclear envelope (**Fig. 4E**, **4F**). Additionally, localization in the plasma membrane demonstrates that this short internal region contains the targeting information required for both compartments. Surprisingly, the fragment comprising residues 430–450 also localized to the plasma membrane (**Fig. 4G**), indicating that plasma membrane targeting can be specified independently of the nuclear envelope determinant. Together, these observations suggest that residues 411–430 include the nuclear envelope targeting determinant, whereas plasma membrane localization is mediated by a broader localization region that includes an additional determinant within residues 430–450.

To further delineate the mitochondrial targeting region, additional truncations at the C-terminus were analysed. The resulting fragment encompassing residues 390–430 localized to both the mitochondria and the nuclear envelope (**Fig.4H**, **4I**), indicating that residues 411–430 retain nuclear envelope targeting activity even in the presence of the mitochondrial targeting determinant. Interestingly, extension of this construct by ten amino acids (390–440) abolished nuclear envelope localization while maintaining exclusive mitochondrial localization (**Fig. S1B**), suggesting that residues 431–440 modulate the activity or accessibility of the nuclear envelope targeting determinant. Likewise, the 390–450 fragment while targeted to mitochondria, it failed to localize in the nuclear envelope (**Fig. S1 C**).

To determine whether these constructs localized to the inner mitochondrial membrane or accumulated within the mitochondrial matrix, we performed fluorescence intensity profile analysis across individual mitochondria. The analysis of the C-terminal region (390–471) produced two distinct GFP fluorescence peaks flanking the MitoTracker-labelled mitochondrial matrix, consistent with enrichment of TtCLS at the mitochondrial membrane (**Fig. 3B**). In contrast, the 390–430, 390–440 and 390–450 fragments lacked the two-peak fluorescence profile characteristic of membrane-associated TtCLS (**Fig. 4H, Fig. S1B, S1C**). Instead, each fragment produced a single GFP fluorescence peak that closely overlapped with the mitochondrial matrix signal. Thus, truncation of the distal C-terminal region progressively converted the mitochondrial localization pattern from membrane-associated, two-peaked profiles to a single matrix-associated fluorescence peak. These results suggest that signal required for mitochondrial import resides within residues 390-430.

The matrix accumulation of these constructs provides important mechanistic insight into TtCLS trafficking. Since proteins that undergo lateral insertion from the TIM23 complex into the inner mitochondrial membrane do not enter the matrix, these observations are more consistent with the conservative sorting pathway, in which proteins are first imported into the mitochondrial matrix before being re-targeted to the inner mitochondrial membrane^33,34^. The inability of the 390–430, 390–440 and 390–450 fragments to reach the inner membrane further indicates that the terminal 21 amino acids (451–471) are dispensable for mitochondrial import but are essential for the subsequent targeting of TtCLS from the matrix to the inner mitochondrial membrane.

Collectively, these findings reveal that the C-terminal segment is considerably more complex than a conventional mitochondrial targeting sequence (**Fig. 4K)**. Rather than functioning solely as a mitochondrial targeting signal, it constitutes a multifunctional localization signal in which overlapping targeting determinants direct TtCLS to the mitochondrial inner membrane, nuclear envelope and plasma membrane (**Fig. 4J, K)**.

### 5. Substitutions at predicted phosphorylation sites alter compartment specific localisation

Having established that the C-terminal region functions as a multi-compartment localization signal directing TtCLS to the mitochondria, nuclear envelope and plasma membrane, we next asked how its compartment-specific localization might be regulated. Because reversible phosphorylation is a common mechanism for modulating protein localization, we examined the C-terminal region for potential phosphorylation sites. Using the phosphorylation-site prediction tool-MusiteDeep^35^, we identified three serine residues S434, S468 and S470 as high-confidence candidate phosphorylation sites

To determine whether phosphorylation regulates mitochondrial targeting, phosphomimetic substitutions mimicking the negative charge of phosphorylation and phosphonull substitutions preventing phosphorylation were introduced into the 390–471 localization sequence. Interestingly, the phosphomimetic substitution S434D completely abolished mitochondrial localization, whereas the corresponding phosphonull mutant retained mitochondrial localization comparable to that of the wild-type fragment (**Fig. 5A**). These observations suggest that phosphomimetic substitution at S434 negatively regulates mitochondrial targeting.

**Figure 5:**
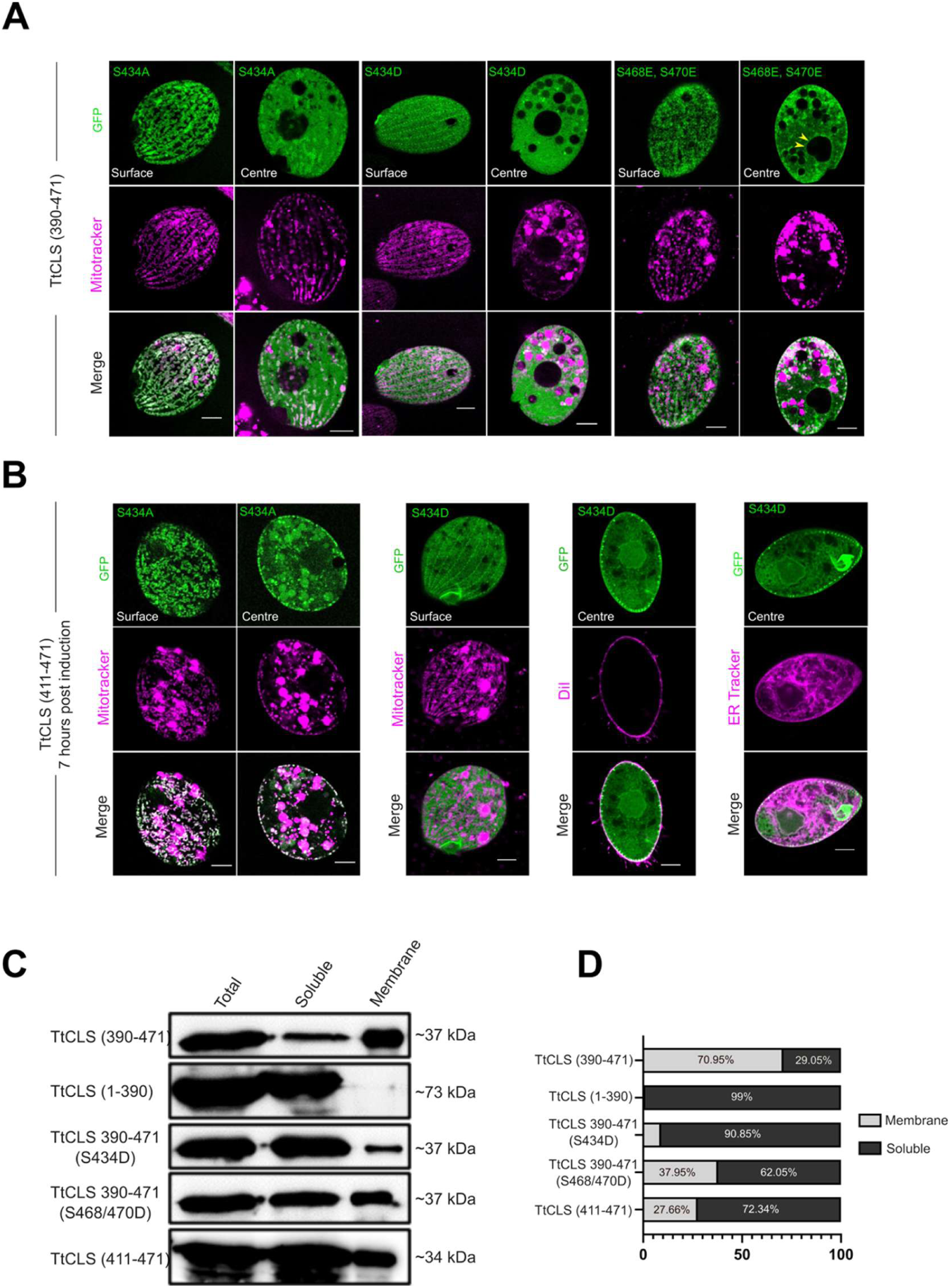
Phosphorylation alters the distribution of TtCLS between cellular compartment. **(A,B)** Representative confocal images of the indicated TtCLS phosphosite mutants of *Tetrahymena* cells from n= 3 independent experiments, >30 cells per experiment, >20 mitchondria per cell, >5 regions of plasma membrane or nuclear envelope labelled with MitoTracker, ER Tracker or DiI as indicated. (**C)** Distribution of the indicated TtCLS constructs between soluble and membrane-enriched fractions assessed by western blotting and its quantification. **(D)** Summary of the localization phenotypes of the indicated phosphosite mutants. Yellow arrow heads indicates the nuclear envelope. Scale bars, 10 μm

We next asked whether the remaining phosphorylation sites function similarly. Simultaneous phosphomimetic substitutions S468E and S470E did not abolish mitochondrial localization but markedly reduced its extent compared with the wild-type localization signal (**Fig. 5A**). Interestingly, this mutant additionally acquired localization to the nuclear envelope, indicating that phospho-mimic substitution at the distal serine residues does not simply inhibit mitochondrial targeting but instead redistributes TtCLS between mitochondria and the nuclear envelope. Together, these observations suggest that phosphorylation potentially fine-tunes the intracellular partitioning of TtCLS rather than functioning as a simple on/off switch for mitochondrial targeting.

The phosphorylation mediated regulation led us to revisit the localization behaviour of the 411–471 fragment, which normally localizes exclusively to the plasma membrane and nuclear envelope. Surprisingly, phosphonull mutations at either S434 or S468/S470 redirected this fragment to the mitochondria, accompanied by a complete loss of plasma membrane and nuclear envelope localization (**Fig. 5B, Fig. S1D**). Careful examination of these cells revealed that the redirected protein accumulated within the mitochondrial matrix rather than the inner mitochondrial membrane. Collectively, these findings indicate that phospho-mimic substitution suppresses the latent mitochondrial targeting activity encoded within the 411–471 region while simultaneously promoting localization to the plasma membrane and nuclear envelope. Phosphorylation thus likely dynamically modulates the activity of overlapping targeting determinants, thereby governing the partitioning of cardiolipin synthase between the mitochondria, nuclear envelope and plasma membrane.

Beyond compartment targeting, the phosphonull analysis also provided insight into mitochondrial membrane association. The accumulation of phosphonull 411–471 mutants within the mitochondrial matrix suggested that residues 390–410, though dispensable for mitochondrial targeting, contribute to association with the inner mitochondrial membrane. To investigate this possibility, membrane association of the localization signal and its derivatives was examined by biochemical fractionation. The 390–471 localization sequence partitioned predominantly into the membrane-enriched fraction (∼71%), supporting its role in membrane association, whereas the remaining catalytic core (1–390) was recovered exclusively in the soluble fraction (**Fig. 5C**, **5D**). Removal of residues 390–410 from the C-terminal region markedly reduced recovery in the membrane fraction, demonstrating that this region is required for efficient membrane binding **(Fig. 5C, D)**.

To identify structural features and potential membrane-interacting elements within the TtCLS C-terminal region, we analysed the protein using AlphaFold 3^36^, PMIPred^37^ and pLMMoRF^37^. Alphafold 3 predicted an α-helix spanning residues 395–416. PMIPred identified an overlapping segment at residues 393-406 as a putative membrane-binding region under both negatively charged and neutral membrane conditions **(Table S2)**. Analysis of the isolated 393-406 peptide also supported membrane interaction with a charge-adjusted free-energy score (ΔΔF_adj) of −20.4 kJ/mol, well below the membrane-binding threshold of −10.0 kJ/mol. Helical wheel analysis suggested amphipathic character (hydrophobic moment of 0.455) **(Table S3, Fig. S4A, S4B)**. Independently, pLMMoRF identified an overlapping membrane-interacting region spanning residues 395-418, with prediction probability exceeding the 0.71 threshold **(Table S2)**. Structure-based analysis using PPM 3.0 with an inner mitochondrial membrane model further predicted a peripheral membrane orientation, with shallow penetration involving residues within the 392–410 region **(Table S3)**. This predicted membrane-contacting surface overlaps the α-helix at residues 395–416 and lies entirely within the region whose deletion reduced membrane association (390-410). Under negatively charged membrane conditions, PMIPred additionally classified residues 416-471 as a potential curvature-sensing region **(Table S2)**.

Taken together, cell fractionation partitioning and modelling strongly and congruently suggest that the 390-410 region, overlaping a predicted amphipathic helix, mediates peripheral attachment to the mitochondrial inner membrane.

### 6. The non-canonical mitochondrial targeting signal of TtCLS is recognized by the conserved mitochondrial import machinery and functions in human cells

Since TtCLS is targeted to the mitochondrial inner membrane through a non-canonical C-terminal localization signal, we asked how this unconventional targeting signal is recognized by the mitochondrial protein import machinery. To address this question, proteins associated with TtCLS were identified by affinity purification followed by mass spectrometric analysis. Functional annotation of the TtCLS interactome revealed multiple components of the mitochondrial protein import machinery, including Pam16/Tim16, a core component of the TIM23 translocase complex, and the small TIM chaperone Tim10, together with several inner mitochondrial membrane and matrix proteins (**Fig. 6A**). The recovery of these proteins indicates that, despite lacking a canonical N-terminal mitochondrial targeting sequence, TtCLS is associated with the conserved mitochondrial protein import machinery.

**Figure 6:**
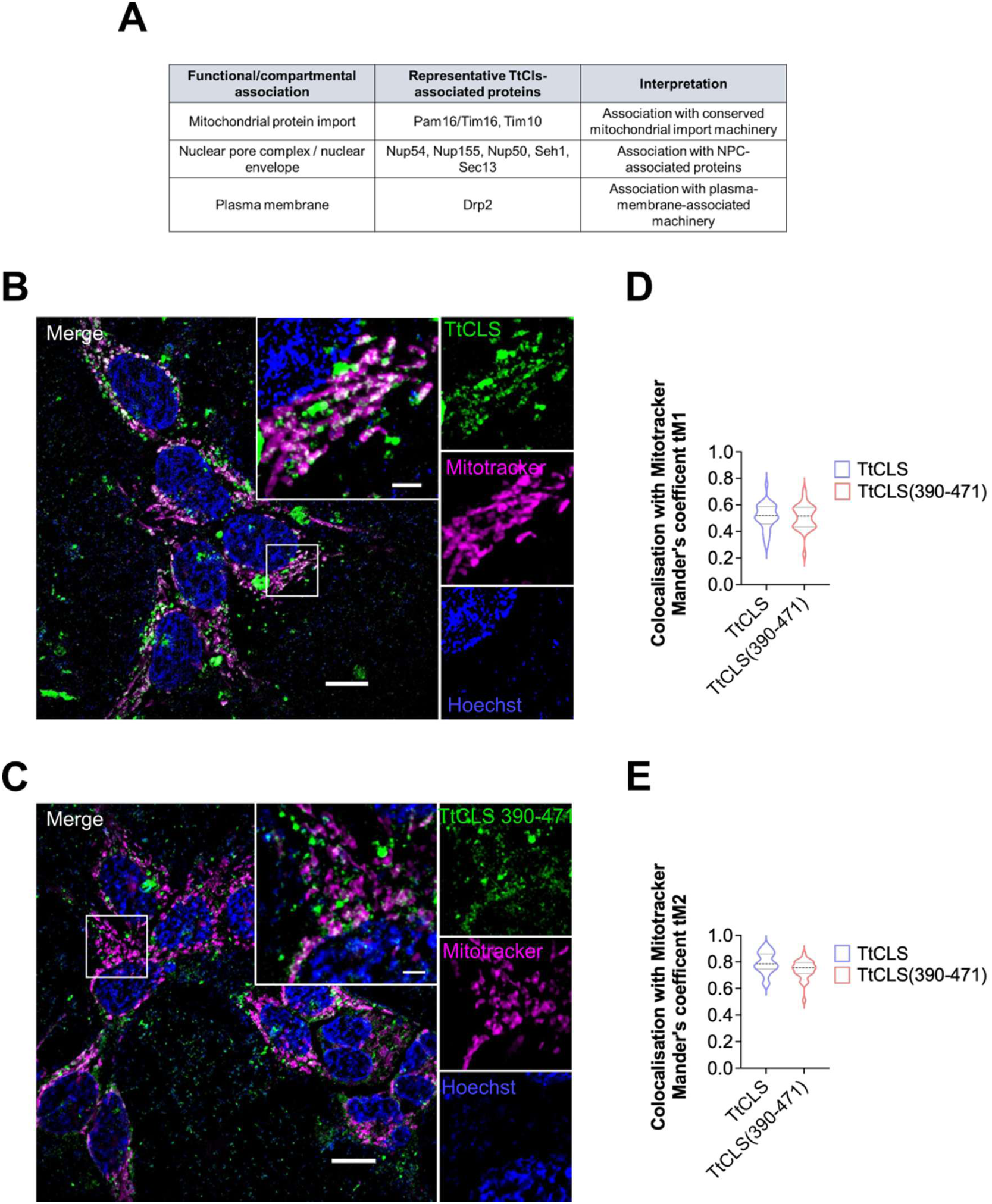
The non-canonical C-terminal signal engages conserved mitochondrial import machinery across eukaryotes. **(A)** TtCLS-associated proteins identified by proteomic analysis following TAP purification of the TtCLS, grouped according to functional or subcellular association. **(B, C)** Representative SR-SIM^2^ images of HEK293 cells expressing GFP–TtCLS (B) or GFP-TtCLS (390–471) (C). Mitochondria were labelled with MitoTracker DeepRed. Both constructs show mitochondrial localisation. Scale bar, 10 μm, Inset Scale bar-1 μm. **(D, E)** Quantification of mitochondrial localisation using Mander’s coefficients. (D) tM1, representing the fraction of GFP signal overlapping MitoTracker Signal. (E) tM2, representing fraction of MitoTracker signal overlapping with GFP signal. Full length TtCLS and TtCLS (390-471) were analysed across more than 30 cells from three independent experiments.

Our truncation analysis unexpectedly suggested that TtCLS follows the conservative sorting pathway employed by a subset of inner mitochondrial membrane proteins. Deletion of the terminal 21 amino acids (451–471) resulted in accumulation of the protein within the mitochondrial matrix rather than the inner mitochondrial membrane (**Fig. S1C**), demonstrating that mitochondrial import occurs independently of these residues. Since proteins that are inserted directly into the inner membrane from the TIM23 complex do not access the mitochondrial matrix^34^, accumulation of the truncated construct within the matrix supports the model in which TtCLS is first imported into the mitochondrial matrix before being subsequently re-targeted to the inner mitochondrial membrane. The identification of Pam16/Tim16 in the TtCLS interactome provides independent biochemical support for this model and suggests that the non-canonical localization signal engages the conserved TIM23-dependent import pathway despite lacking a canonical N-terminal presequence.

The ability of this localization signal to engage the conserved import machinery prompted us to ask whether it is recognized exclusively in ciliates or whether it can function in evolutionarily distant eukaryotes. To address this, both GFP-tagged TtCLS and GFP-tagged TtCLS (390-471) fragment were independently expressed in HEK293 cells and its intracellular localization was examined by SR-SIM^2^ (**Fig. 6B**, **6C**). Both proteins localised to mitochondria and colocalised with MitoTracker. Manders’ colocalization analysis quantified the fraction of GFP signal overlapping with MitoTracker (tM1) and the fraction of MitoTracker signal overlapping with GFP (tM2) (**Fig. 6D**, **6E**). Full-length TtCLS exhibited tM1 and tM2 coefficients of (0.5112 ± 0.097) and (0.7925 ± 0.080), respectively, while fragment TtCLS (390–471) exhibited corresponding values of (0.5080± 0.096) and (0.7411 ± 0.070) (**Fig. 6D**, **6E**). This analysis included more than 30 cells across three independent experiments. Values are presented as mean ± standard deviation (SD). These findings demonstrate that C-terminal region is sufficient to promote mitochondrial localisation in human cells, indicating that targeting information is recognisable by the mitochondrial targeting machinery of evolutionarily distant eukaryotes, despite the absence of canonical N-terminal mitochondrial targeting peptide.

Collectively, these findings demonstrate that the C-terminal localization signal is compatible with the evolutionarily conserved TOM/TIM protein import machinery. Together with the matrix accumulation of the 390–450 truncation mutant **(Fig. S1B)**, our results support a model in which TtCLS is first imported into the mitochondrial matrix before being re-targeted to the inner mitochondrial membrane through the conservative sorting pathway. More broadly, these observations reveal that mitochondrial protein import can be achieved through a structurally distinct, C-terminal localization signal that is nevertheless recognized by the canonical mitochondrial import machinery across evolutionarily distant eukaryotes.

### 7. Conservation and evolution of CLS_PLD cardiolipin synthases in bacteria and eukaryotes

To determine whether features of the TtCLS C-terminal region are unique to *Tetrahymena* or conserved more broadly, we compared TtCLS with 162 homologues, representing 163 CLS sequences from 156 bacterial and eukaryotic taxa. The catalytic core of CLS_PLD was highly conserved across the dataset but contained lineage-specific signatures, including several characteristic charged residues within the conserved motifs of alveolate sequences (**Fig. S2**). In contrast, the lengths of both terminal regions differed markedly among groups. We therefore reconstructed the phylogeny of CLS_PLD to interpret the evolution of these regions (**Fig. 7**).

**Figure 7:**
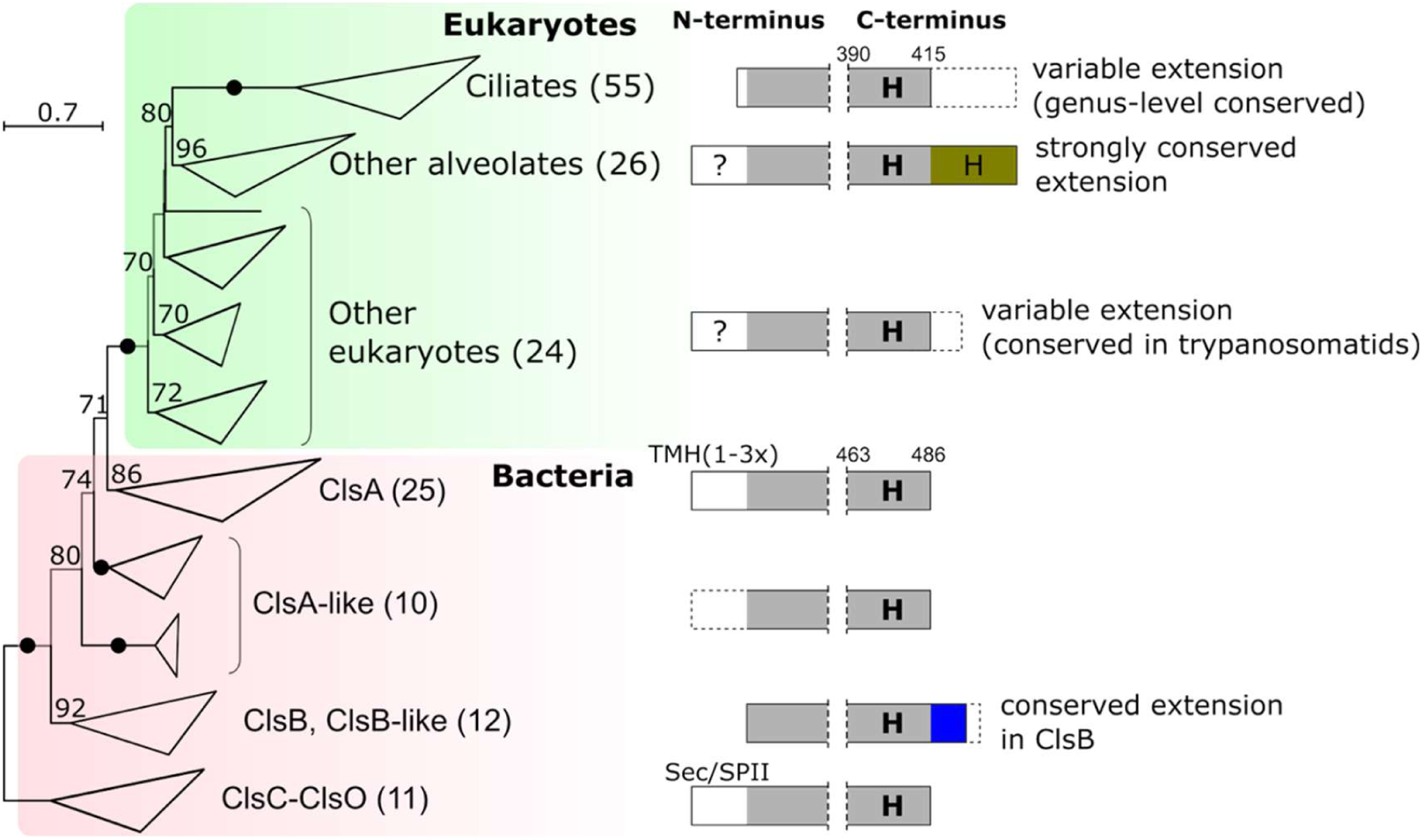
Phylogeny and terminal-region architecture of bacterial and eukaryotic CLS_PLD cardiolipin synthases. Maximum-likelihood phylogeny of 163 sequences inferred using IQ-TREE under the LG+I+R6 model, with selected clades collapsed. Numbers in parentheses indicate sequence counts. Ultrafast bootstrap support values ≥70 are shown at branches; filled black circles indicate support >99. Generalized protein architectures for selected groups are shown on the right, with residue positions numbered according to TtCLS and *E.coli* ClsA. Grey boxes represent aligned regions, with the intervening sequence omitted at the dashed vertical lines. Conserved C-terminal extensions are coloured. Dashed-outline boxes indicate regions variably present within each group. **H**, predicted α-helix (the conserved amphipathic helix is shown in bold); TMH, transmembrane helix; Sec/SPII, Sec signal peptide cleaved by signal peptidase II. The scale bar indicates substitutions per site. The fully expanded tree and targeting predictions are shown in **Figure S3.**

An α-helix corresponding to the proposed membrane-associating helix of TtCLS was predicted in all ten representative bacterial and eukaryotic CLS proteins examined using AlphaFold 3, despite weak conservation of its primary sequence. The conservation of this helix raises the possibility that it contributes to membrane association across CLS_PLD proteins, even in the absence of a downstream extension. Indeed, most bacterial CLS proteins terminated immediately after this helix, as exemplified by *Escherichia coli* ClsA. However, a clade of nine bacterial sequences containing *E. coli* ClsB possessed a partially conserved C-terminal extension averaging 34 residues. These extensions were enriched in alanine and proline and depleted in aromatic residues relative to the mature catalytic core (**Table S4**). ClsB lacks N-terminal transmembrane domains characteristic of ClsA or Sec/SPII signals characteristic of ClsC/ClsO, so the presence of conserved C-terminal extensions in them is notable. When the tree was rooted with the more divergent ClsC/ClsO clade, the eukaryotic sequences were most closely related to the bacterial ClsA clade than to ClsB, consistent with an independent acquisition of the ClsB C-terminal extensions **(Fig. 7).**

Eukaryotic sequences varied considerably at both termini. Only three of the 105 eukaryotic sequences contained a predicted canonical N-terminal mitochondrial targeting sequence (**Fig. S3**). C-terminal extensions beyond the ClsA terminus were generally short and poorly conserved, or absent in eukaryotes other than alveolates; only trypanosomatids had a conserved, negatively charged extension of approximately 20 residues. Strikingly longer C-terminal extensions were present in alveolate CLSs. Those in apicomplexans, dinoflagellates and, notably, the deep-branching alveolate *Colponema* were strongly conserved, hydrophobic and leucine-rich. The region beyond the ClsA terminus had an average length of 43 residues and was consistently predicted to contain an additional α-helix. Ciliate proteins were also commonly C-terminally extended, but their extensions were generally conserved only within genera and varied substantially at deeper taxonomic levels. Ciliate sequences also exhibited markedly greater sequence divergence than the other sequences in the dataset **(Fig. 7).** The extensions of *Tetrahymena* and *Paramecium* averaged 56 and 28 residues, respectively, beyond the conserved ClsA terminus **(Table S4).**

Given the apparent functional roles of the TtCLS C-terminal region, we next tested whether it is evolutionarily constrained despite its sequence divergence from other ciliates. Codon-wise FEL analysis of 12 CLS sequences from the Tetrahymena lineage revealed pervasive purifying selection across much of the protein, including the C-terminal region, but no pervasive positive selection (Materials and Methods). Regional CODEML estimates likewise showed comparable dN/dS in the C-terminal and main protein regions (ω = 0.0114–0.0138 versus 0.0125–0.0127). Excluding the CLS_PLD catalytic motifs had negligible effect, whereas the motifs themselves showed even stronger constraint (ω ≈ 0.0065), as expected. Thus, the C-terminal region is subject to strong purifying selection comparable to that of the main protein.

To investigate whether membrane-interacting C-terminal features extend beyond *Tetrahymena*, we analysed the *Plasmodium falciparum* CLS (PfCLS) using PMIPred, pLMMoRF and PPM 3.0. PMIPred identified a candidate membrane-binding region spanning residues 543-561 (RNFLQIFMSYCAYHLGKLS), with the isolated peptide yielding a charge-adjusted free energy score (ΔΔF_adj) of −18.872 kJ/mol **(Table S2)**. The peptide had a net charge of +2 and a hydrophobicity value of 0.669 **(Table S2)**. Independently, pLMMoRF identified a C-terminal region spanning residues 545-601 with prediction probabilities above the 0.71 cutoff **(Fig. S4, Table S2)**. PPM 3.0 likewise predicted 11 membrane-embedded residues within the C-terminal region, spanning positions 545–603 together with four at positions 173–178 and three at positions 267–273. Together, these analyses support membrane-interaction potential in the PfCLS C-terminal region, which includes both the α-helix corresponding to the membrane-associating helix of TtCLS and the additional predicted downstream α-helix (**Fig. 7**). Because both TtCLS and PfCLS are predicted to contain a homologous membrane-interacting helix, and this helix is also conserved in the other CLS_PLD sequences we examined, we tested its membrane-binding potential with PPM 3.0 in an additional four bacterial and six eukaryotic sequences **(Table S3).** Strikingly, in all twelve sequences the predicted membrane-contacting residues overlapped this helix, including bacterial ClsA sequences that are predicted to have N-terminal transmembrane helices. These results suggest that CLS_PLD enzymes across domains carry a potential for peripheral membrane association.

## Discussion

Protein sorting is in general mediated by a N-terminal signal sequence^38^. However, C-terminal and internal signals are also known to mediate targeting of proteins to different compartments including mitochondria^39–41^. For example, while BCS1 and cytochrome c1 use an internal signal^40,42^, a C-terminal determinant targets cytochrome b5 and VAMP-1B for their mitochondrial import^39,43^. Although a single signal sequence targeting to two compartments is known, targeting to three different compartments is unusual. The present study reports that a single protein, TtCLS localises to three independent compartments **(Fig. 1)** and the targeting determinants for all three compartments reside as an overlapping sequence within a lineage specific C-terminal extension **(Fig. 4),** making it a novel mechanism for multi-compartment targeting.

The differential localisation of a protein is generally regulated by post-translational modifications such as phosphorylation^3^. As for example phosphomimetic substitutions re-distributes a tumour suppressor protein PTEN from plasma membrane to cytosol thereby diminishing its membrane dependent phosphatase activity^44^. Similarly, phosphorylation also induces distribution of a protein in MAP kinase pathway K-Ras4B from plasma membrane and other intracellular membranes including endoplasmic reticulum and outer mitochondrial membrane thereby inducing cellular death^45^. By using phosphomimetic and phospho-defective mutants we demonstrate that the localisation of a bacterial type cardiolipin synthase in *Tetrahymena* can also be regulated by phosphorylation **(Fig. 5A**, **5B)**. Intriguingly, phosphosite mutation also affects membrane binding affinity of TtCLS **(Fig. 5C**, **5D)**, a finding also reported in PTEN where phosphorylation at its C-terminal tail inhibits membrane binding^44^. Overall these results suggest presence of a conserved mechanism involving phosphorylation for regulating differential distribution coupled with membrane binding across evolutionarily distant organisms. Interestingly, our finding that the fragment containing residues 411-430 has the information for targeting to all the three compartments, but it requires either presence of an additional 21 residues at its N-terminus (390-410) or rest of the C-terminal residues (431-471) with dephosphorylated state at one or more phosphosites for targeting to mitochondria (**Fig. 4E, 4F, 4H, 5B, S1D**). This is a novel example of protein targeting where in addition to phosphorylation, the surrounding sequence also regulates dynamic distribution of a protein among various cellular compartments.

The mechanism by which this unconventional targeting region directs TtCLS to the mitochondrial inner membrane is equally intriguing. Deletion of the last 21 amino acids resulted in matrix accumulation **(Fig. S1)**, suggesting that TtCLS passes through the mitochondrial matrix before being sorted to the inner membrane. Consistent with this model, the TtCLS interactome identified conserved mitochondrial import components, including Pam16/Tim16 and Tim10 **(Fig. 6A)**. Furthermore, expression in mammalian cells demonstrated that the *Tetrahymena* targeting region is recognized by mitochondrial import machinery in an evolutionarily distant host **(Fig. 6B – 6E)**. This cross-lineage compatibility raises the possibility that C-terminal mitochondrial targeting signals share common recognition determinants across eukaryotes.

One of the most striking findings of this study is that the mitochondrial targeting information of TtCLS resides entirely within its C-terminal region (**Fig. 3**, **Fig. 4)**. TtCLS belongs to the bacterial phospholipase D-type cardiolipin synthase family (CLS_PLD), which is retained in a limited number of eukaryotic lineages, including ciliates and apicomplexans^19^. Although its catalytic core is highly conserved across bacterial and eukaryotic homologues, its C-terminal region is markedly extended relative to bacterial and several other eukaryotic CLS_PLD proteins **(Fig. S2)**. Our deletion analysis demonstrates that this extension is necessary and sufficient for mitochondrial localization, suggesting that its evolutionary acquisition may have introduced novel targeting information (**Fig. 3)**. Notably, other alveolate CLS proteins also possess C-terminal extensions of varying lengths, raising the possibility that these regions contribute to their intracellular targeting.

The membrane association of TtCLS residues 390–410 (**Fig 5C**, **5D**) may be conserved across the CLS_PLD family **(Table S3)**. Computational analyses predict membrane-association potential within α-helices at homologous positions in representative bacterial and eukaryotic CLS_PLD proteins, while predicted transmembrane domains are absent from our 163-sequence dataset except in the N-terminal regions of ClsA homologues **(Fig. 7)**. Despite possessing these N-terminal hydrophobic segments, *B. subtilis* ClsA associates with septal membranes through two C-terminal amphipathic α-helices, either of which is sufficient for septal membrane localization^46^. Similarly, *E. coli* ClsA remains membrane-associated and catalytically active following deletion of its N-terminal hydrophobic segments^47^. *E. coli* ClsB and ClsC lack predicted transmembrane helices altogether, yet their activity is recovered in membrane fractions^48^. Among eukaryotes, we could not confirm the two C-terminal transmembrane domains proposed for *Trypanosoma brucei* CLS^49^; instead, we predict a C-terminal amphipathic helix and additional potential membrane-contact sites nearer the N-terminus (Table S4). Similarly, PfCLS contains the conserved helix alongside other predicted membrane-interacting regions, including the downstream helix conserved among non-ciliate alveolates and a region overlapping the hydrophobic segment at residues 163–185 reported by Adhur et al. (2026)^25^. This suggests a previously unrecognized, potentially ancestral mechanism of membrane association through a C-terminal amphipathic helix, shared by bacterial and eukaryotic CLS_PLD enzymes. The multifunctional targeting architecture of TtCLS may therefore have evolved around this ancestral membrane-binding element.

The extra-mitochondrial localization of TtCLS raises questions about its physiological significance. Its presence at the nuclear envelope is consistent with previous reports of cardiolipin at this membrane^23^ and its involvement in nuclear expansion^24^. Similarly, plasma-membrane localization points to another potential site of extra-mitochondrial cardiolipin metabolism. Notably, our recent findings identify a role for the plasma-membrane cardiolipin pool in rapid nuclear expansion (S. Rout, S. Sahoo and A. Rahaman, 2026, data not shown). A broader parallel emerges from *P. falciparum* CLS_PLD, which localizes predominantly to mitochondria in asexual blood stages but also accumulates at the cellular periphery in gametocytes, coinciding with cardiolipin detection at the plasma membrane^25^. Together, these observations suggest that, unlike the predominantly mitochondrial, N-terminally targeted CLS_CAP enzymes, CLS_PLD proteins may confer greater spatial and developmental flexibility in the regulation of cardiolipin pools, potentially mediated by C-terminal targeting determinants such as those identified here in TtCLS.

In conclusion, our study identifies an unconventional protein-targeting strategy in which a single C-terminal region integrates overlapping determinants for localization to three membrane compartments, with their relative contributions potentially modulated by phosphorylation. Although TtCLS lacks a canonical N-terminal mitochondrial targeting sequence, its C-terminal region can direct mitochondrial localization in mammalian cells, suggesting that recognition of C-terminal targeting information may be evolutionarily conserved across eukaryotes. The same region also contains an amphipathic helix whose membrane-association function may represent an ancestral feature of the CLS_PLD family. Together, these findings expand current models of mitochondrial protein targeting by revealing an unexpected degree of evolutionary plasticity in targeting architecture, and illustrate how evolution can remodel protein-targeting information to generate new patterns of intracellular membrane distribution while preserving compatibility with conserved cellular transport systems.

## Materials and Methods

### 1. *Tetrahymena* strains and culture conditions

*Tetrahymena thermophila* strains CU428 and B2086 were obtained from the *Tetrahymena* Stock Center (Cornell University, Ithaca, NY, USA). Cells were routinely cultured in Super Proteose Peptone (SPP) medium containing 2% (w/v) proteose peptone (Gibco, 211684), 0.2% (w/v) glucose (Sigma-Aldrich, G7021), 0.1% (w/v) yeast extract (Gibco, 212750), and 0.003% (w/v) ferric EDTA (Sigma-Aldrich, 03650) supplemented with 1× antibiotic–antimycotic solution (penicillin, streptomycin and amphotericin B; HiMedia, A002A). Cultures were maintained at 30°C with shaking at 90 rpm and harvested during logarithmic growth phase (2 × 10⁵ to 4 × 10⁵ cells mL⁻¹) unless otherwise stated. Cell density was determined using a haemocytometer.

For starvation experiments, cells were washed once with and resuspended in DMC medium consisting of 0.17 mM sodium citrate, 0.1 mM NaH₂PO₄, 0.1 mM Na₂HPO₄, 0.65 mM CaCl₂ and 0.1 mM MgCl₂. Unless otherwise indicated, both live and fixed cells were harvested by centrifugation at 1,100 × g for 2 min.

### 2. Generation of TtCLS truncation and phosphomutant constructs

A series of N- and C-terminal truncation constructs of *Tetrahymena thermophila* cardiolipin synthase (TtCLS) were made. C-terminal truncations were generated by introducing stop codons (TGA) at the desired positions using site-directed mutagenesis (QuikChange; Agilent Technologies). N-terminal truncations were generated by PCR amplification of the corresponding regions using specific primers, followed by directional cloning into pENTR/D-TOPO entry vector (Invitrogen, K240020). Entry clones were recombined into the destination vector, pIGF using Gateway LR Clonase II (Invitrogen, 11791020) to generate constructs for expressing N-terminal GFP fusion proteins.

Phosphomimetic (S434D, S468E/S470E, and S434D/S468E/S470E) and phosphonull (S434A, S468A/S470A, and S434A/S468A/S470A) mutants were generated by site-directed mutagenesis using pENTR/D constructs as template, verified by sequencing and recombined into destination vector (pIGF) by LR Clonase reaction.

### 3. Generation of constructs for mitochondrial labelling

Constructs for various mitochondrial translocons were generated to identify the sub-mitochondrial localization of TtCLS. The TtOXA1 protein, previously identified as mitochondrial protein^32^, was PCR-amplified from *Tetrahymena thermophila* genomic DNA using gene-specific primers. The amplified products were cloned into the pVGF expression vector at the XhoI and ApaI restriction sites to generate constructs encoding N-terminal GFP tagged TtOXA1.

### 4. Transformation of *Tetrahymena* cells using electroporation

For electroporation, 50 mL cultures of the complementary mating types CU428 and B2086 were grown to a density of approximately 4 × 10⁵ cells mL⁻¹, harvested by centrifugation, and resuspended in DMC starvation medium at a final density of 2.5 × 10⁵ cells mL⁻¹. Following 16–18 h of starvation, equal volumes of the two mating types were mixed to initiate conjugation. Approximately 20 μg of plasmid DNA was introduced into conjugating cells by electroporation using a BTX Gemini X^2^ electroporation system.

Following electroporation, cells were allowed to recover overnight in SPP medium before selection with 100 μg mL⁻¹ paromomycin sulfate. Stable transformants were maintained under selection, and expression of the GFP-tagged constructs was induced by supplementing SPP medium with 0.5–2 μg mL⁻¹ CdCl₂ for 3–6 h.

### 5. Biolistic transformation

Co-expression strains were generated by biolistic transformation using the Bio-Rad PDS-1000/He particle delivery system, following a previously described protocol^50^. Briefly, *Tetrahymena thermophila* cells expressing GFP-tagged TtOXA1 under paromomycin selection were starved for approximately 16 h and subsequently transformed with a construct encoding mCherry-tagged TtCLS. The mCherry-TtCLS construct was targeted to MTT1 locus by homologous recombination.

Transformants were selected in SPP medium supplemented with 60 μg mL⁻¹ blasticidin and 1 μg mL⁻¹ CdCl₂. Following isolation of stable transformants, cells were maintained under blasticidin selection in the presence of CdCl₂. For fluorescence imaging, transformants were grown without blasticidin and cadmium chloride, and transgene expression was induced by adding 1 μg mL⁻¹ CdCl₂ for 3–5 h before imaging.

### 6. Tetrahymena cell fixation

*Tetrahymena thermophila* cells expressing fluorescently tagged proteins were grown in SPP medium to a density of 2.0–4.0 × 10^5^ cells mL⁻¹. Expression of transgenes under the control of the cadmium-inducible MTT1 promoter was induced by supplementing the culture medium with 0.5 μg mL⁻¹ CdCl₂ for 3–5 h. Cells were harvested by centrifugation at 1,100 × g for 2 min and fixed for 20 min in 4% (w/v) paraformaldehyde (PFA) prepared in 50 mM HEPES, pH 7.5. For experiments requiring optimal preservation of mitochondrial architecture, including super-resolution structured illumination microscopy (SR-SIM²), fixation was performed for 1 h at 4°C.

### 7. Membrane and organelle staining

For plasma membrane labeling, 2 mM DiI (1,1′-Dioctadecyl-3,3,3′,3′-tetramethylindocarbocyanine perchlorate) (Sigma-Aldrich, 42364) was prepared in ethanol and used at a final concentration of 5 μM. Fixed cells were incubated with the dye for 10 min at room temperature and washed twice with 10 mM HEPES, pH 7.5 before imaging.

MitoTracker Deep Red FM (Invitrogen, M22426) for labelling mitochondria and ER-Tracker™ Red (Invitrogen, E34250) for endoplasmic reticulum were prepared in DMSO and used at a final concentration of 0.5 μM. Cells were incubated with the dye for 1 h at 30°C with shaking at 90 rpm and fixed before imaging. Live cell imaging was also performed while imaging MitoTracker labelled mitochondria.

### 8. Gene synthesis and codon optimisation

The full length TtCLS coding sequence was codon-optimised for expression in heterologous eukaryotic systems like mammalian cells, to account for the differences in genetic codon usage between *Tetrahymena* and other heterologous expression hosts. In ciliates like *Tetrahymena*, UAA and UAG encode for glutamine, whereas UGA functions as the only stop codon. The degeneracy of the genetic code was used to preserve encoded amino acid sequence while replacing codons. The codon optimised gene was synthesised commercially by Barcode Biosciences Pvt. Ltd. (India) and used for subsequent cloning and expression experiments.

### 9. HEK cell culturing and transfection

HEK293 cells were routinely cultured in DMEM media supplemented with 10% fetal bovine serum (FBS). Cells were seeded onto circular cover glass placed in 12-well sterile culture dishes. Cells upon reaching a confluency of 40% were transfected with 300 ng plasmid per well using Lipofectamine LTX (Thermo Fisher Scientific) according to manufacturer’s instructions. DNA-lipid complex was prepared and incubated at room temperature for 25 minutes before adding it to the cells. Following transfection, cells were allowed to recover for 48 h in standard culture conditions. Mitochondria were labelled with 50nM MitoTracker Deep Red FM for 30 minutes, followed by staining with 1 μg mL⁻¹ Hoechst to label the nuclei. Cells were fixed with 4% paraformaldehyde (PFA) and were subjected to super-resolution structured illumination microscopy (SR-SIM^2^).

### 10. Confocal microscopy

Confocal fluorescence imaging was performed using either a Leica SP8 DMi8 or a Zeiss LSM 780 laser-scanning confocal microscope equipped with a 63×/1.4 NA Apochromatic oil-immersion objective and 405/488/552/638 nm lasers. Images were acquired using Leica Application Suit X (LAS-X) or Zen Black and analysed using FIJI/ImageJ (NIH, USA). The number of independent experiments and number of cells analysed are indicated in the figure legend. Coloc2 was used to access the Manders’ colocalization.

### 11. Super-resolution Structured illumination microscopy (SR-SIM²)

Super-resolution structured illumination microscopy (SR-SIM²) imaging was performed using a Zeiss Elyra 7 microscope equipped with 63×/1.4 NA Apochromatic oil-immersion objective. The image reconstruction was carried out using Zeiss ZEN Black application and processing was performed using Zeiss Zen Blue application.

### 12. Fluorescence intensity profile analysis

Fluorescence intensity profiles were generated using the Plot Profile function in FIJI/ImageJ. Straight line regions of interest were drawn across individual mitochondria, plasma membrane and nuclear envelope. The fluorescence intensities corresponding to GFP and the respective organelle marker were extracted simultaneously for a region of interest. The extracted fluorescence values were plotted using GraphPad Prism 8.

For analysis of sub-mitochondrial localization, the relative positions of fluorescence intensity peaks were compared. Coincident fluorescence peaks were interpreted as colocalization, whereas spatially separated peaks were used to distinguish sub-mitochondrial localization.

### 13. Live-cell immobilization

*Tetrahymena* cells (2–6 × 10⁶ cells mL⁻¹) were immobilized by mechanical confinement. Briefly, 4 μL of cells was mounted on a glass microscope slide and immediately overlaid with an 18 × 18 mm No. 1 coverslip supported at the four corners with petroleum jelly. This preparation effectively restricted cell movement for approximately 30 min, providing a sufficient imaging window for live-cell fluorescence microscopy.

### 14. Affinity purification of TtCLS protein complexes for interactome analysis

*Tetrahymena* cells expressing TAP-tagged TtCLS were cultured in 4 L SPP medium to ∼2.5 × 10⁵ cells mL⁻¹ and induced with 2 μg mL⁻¹ CdCl₂ for 5 h at 30°C and 90 rpm. Cells were harvested at 1,100 × g for 4 min, lysed in ice-cold buffer containing 20 mM Tris-HCl, pH 8.0, 100 mM NaCl, 2 mM MgCl₂, 0.2 mM EGTA, 0.1% Tween-20, 10% glycerol, 1 mM DTT, and protease inhibitors, and clarified by ultracentrifugation at 250,000 × g for 1 h at 4°C. The clarified lysate was incubated with rabbit IgG agarose resin for 2 h at 4°C, and then subjected to TEV cleavage. Eluted proteins were analysed by SDS-PAGE and silver staining before mass spectrometric identification of associated proteins.

### 15. Membrane fractionation

*Tetrahymena* cells (∼1.5 × 10^6^ cells) expressing various GFP tagged proteins were resuspended in 1 mL ice-cold buffer containing 25 mM HEPES (pH 7.4), 150 mM NaCl, 10% glycerol, and 1 mM EDTA, supplemented with protease inhibitor cocktail (Roche). Cells were passed through a clearance gap of 18 μm using a ball-bearing homogenizer (isobiotec). The intact cells and cellular debris were removed by centrifugation at 1,000 × g for 10 min at 4°C, and the resulting supernatant was further centrifuged at 21,000 × g for 30 min at 4°C. The supernatant was collected as the soluble fraction and pellet as the membrane-enriched fraction. The fractions were analysed by western blotting.

### 16. Western blot analysis

For western blotting, rabbit anti-GFP antibody (Invitrogen, A11122; 1: 4,000) was used as the primary antibody, followed by HRP-conjugated anti-rabbit IgG antibody (AbClonal, AS014; 1: 10,000) as the secondary antibody. Signal production was initiated by adding enhanced chemiluminescence solution (ECL; Bio-Rad, 170-5061) and signal was detected using the BioRad ChemiDoc Imaging system.

### 17. Sequence analysis

The TtCLS protein sequence (gene identifier: TTHERM_01109760) was retrieved from the *Tetrahymena* Genome Database (https://tet.ciliate.org). Subcellular localization and targeting signals were predicted using TargetP 2.0^27^, SignalP 6.0^28^, DeepLoc 2.0^29^, DeepMito^30^ and WoLF PSORT^31^. The tertiary structure of TtCLS was modelled using AlphaFold 3 (https://alphafoldserver.com). Membrane association was assessed using PMIPred (https://pmipred.fkt.physik.tu-dortmund.de/curvature-sensing/), which employs a transformer neural network trained on molecular dynamics-derived interaction data to predict protein-membrane interaction energetics and classify regions as non-binding, membrane-binding or curvature-sensing. PMIPred was run using the highest-ranked AlphaFold 3 model with default settings for both negatively charged and neutral membranes. Sequence-based predictions of membrane-interacting molecular recognition features were obtained using pLMMoRF (https://plmmorf.hegelab.org/), which combines protein language model representations with a deep convolutional neural network. Predictions were generated with default settings, applying the default probability threshold of 0.71. Membrane positioning was predicted using PPM 3.0 (https://opm.phar.umich.edu/ppm_server3_cgopm)^51^, with the highest-ranked AlphaFold 3 model, the inner mitochondrial membrane setting, and otherwise default parameters for planar membranes^51^. The analysis was extended to CLS_PLD sequences from *Plasmodium falciparum*, six other eukaryotes and four bacteria (**Table S3**), using top-ranked AlphaFold 3 models and the appropriate membrane setting (inner mitochondrial, inner gram negative, gram positive) for all sequences. The helices were identified in the AlphaFold models, whereas membrane-embedded residues were predicted using PPM 3.0. A helix was considered membrane-interacting when it overlapped the predicted membrane-embedded residues, and its homology to the TtCLS helix (residues 395–416) was verified in a sequence alignment.

The TtCLS protein sequence was used as a query in BLASTP sequence similarity searches to retrieve characterized homologs from UniProtKB/Swiss-Prot and additional representative eukaryotic and bacterial sequences from NCBI nr and tsa_nr databases. Ciliate sequences were retrieved from the *Tetrahymena* Genome Database, ParameciumDB (https://paramecium.i2bc.paris-saclay.fr/), the P10K database (*Colpidium striatum*; https://ngdc.cncb.ac.cn/p10k/), and individual ciliate transcriptomic studies. Predicted splice boundaries in seven *Tetrahymena* species (*T. canadensis*, *T. malaccensis*, *T. shanghaiensis*, *T. pyriformis*, *T. vorax*, *T. utriculariae* and *T. paravorax*), primarily within the N-terminal coding region, were adjusted based on conserved gene structure within the genus. Ciliate transcripts retrieved as contigs were translated using species-specific genetic codes. Redundant, highly similar paralogs and most transcript-derived sequences lacking a terminal stop codon were excluded. Exceptions were retained for the phylogenetically distinct taxa Metopus sp. and Prodiscophrya collini, the latter of which already contained a long C-terminal extension. The final dataset contained 163 CLS_PLD homologs, which were aligned using the L-INS-i strategy implemented in MAFFT v7.525^52^. Poorly aligned regions were removed using BMGE v1.12 with the parameters −m BLOSUM30 −b 3 −g 0.4. Maximum-likelihood phylogenetic inference was performed using IQ-TREE v3.0.1^53^ under the best-fitting substitution model, LG+I+R6, selected by ModelFinder. Branch support was assessed using 1000 ultrafast bootstrap replicates (UFBoot2). N-terminal mitochondrial transit peptides were predicted for eukaryotic sequences using TargetP 2.0. Transmembrane regions throughout the available protein sequences and N-terminal signal peptides were predicted for bacterial and eukaryotic sequences using DeepTMHMM 1.0^57^ and SignalP 6.0^28^, respectively. Some targeting signals were predicted when downstream methionines within the N-terminal extensions were used as sequence starts. Sequence logos of *Tetrahymena* sequences and CLS_PLD catalytic motifs (**Fig. S2**) were generated using Jalview v2.11.5.3 from a reduced version of the full alignment (excluding sequences with gaps in either catalytic region).

Pervasive site-specific selection across coding regions from 12 species of the *Tetrahymena* clade, including *Tetrahymena* spp. and *Colpidium striatum*, was assessed from a codon-based alignment (generated by MAFFT L-INS-i and manually codon-adjusted) using Fixed Effects Likelihood (FEL v2.6) in HyPhy through the Datamonkey web server^54,55^. FEL independently estimates synonymous and nonsynonymous substitution rates at each codon and tests for purifying or diversifying selection using likelihood-ratio tests^54^. Analyses used the ciliate nuclear genetic code, allowed synonymous-rate variation, and employed 1,000 parametric-bootstrap replicates to derive site-level null distributions. Regional selective constraint was assessed on the same alignment using CODEML in PAML v4.10.10^56^ to estimate dN/dS (ω). CODEML analyses used the ciliate nuclear genetic code, F3×4 codon frequencies and the phylogenetic topology inferred by FEL, with gaps retained as missing data. A partitioned model allowing evolutionary parameters to differ between regions (Mgene = 4) provided a better fit than a shared-parameter model (Mgene = 0) and was used to obtain regional ω estimates. *T. thermophila* C-terminal regions 416–471 and 390–471 were compared with the main protein region (1–389) and the CLS_PLD catalytic motifs (142–160, 329–346 and 348–357); sensitivity analyses repeated comparisons after excluding the catalytic motifs from the main region.

### 18. Statistical Analysis

Statistical analysis and graphical representations were performed using GraphPad Prism 8. Quantitative data are presented as mean ± standard deviation. The number of independent experiments and number of cells are mentioned in the corresponding figure legends. P < 0.05 was considered statistically significant.

## Acknowledgements

We thank the BRIC-ILS Mass Spectrometry Facility at the Institute of Life Sciences, Bhubaneswar and Dr Amol Suryawanshi for proteomics analyses. Thanks to Debraj Koiri for his help in performing transfection experiments. We also thank Mr. Sakti Ranjan Rout, Ms. Gargi Dey and Ms. Urbi Ghosh of the AR laboratory for their help and support throughout this study. JJ was supported by the Czech Science Foundation grants 21-19798M and 24-12396S.

## Supplemental information

### Supplementary Figures

- Figures S1. Mitochondrial localization of C-terminal TtCLS fragments in *Tetrahymena thermophila*
- Figure S2. Conservation of the C-terminal region and catalytic motifs of CLS_PLD cardiolipin synthases
- Figure S3. Expanded phylogeny and predicted targeting features of CLS_PLD cardiolipin synthases
- Figure S4: Predicted membrane interaction properties of selected C-terminal peptides from TtCLS and PfCLS using PMIPred

### Tables

- Table S1: In-silico prediction of targeting signals in TtCLS using different tools
- Table S2: PMIPred and pLMMoRF prediction for membrane binding segment in TtCLS and PfCLS
- Table S3: PPM 3.0 prediction of membrane interactions in selected CLS_PLD proteins based on AlphaFold 3 models (MAH, membrane-associated helix homologous to TtCls 395–416)
- Table S4: Amino acid composition of CLS C-terminal extensions and mature cores (excluding N- and C-terminal extensions) in five organismal groups

## List of constructs and reagents

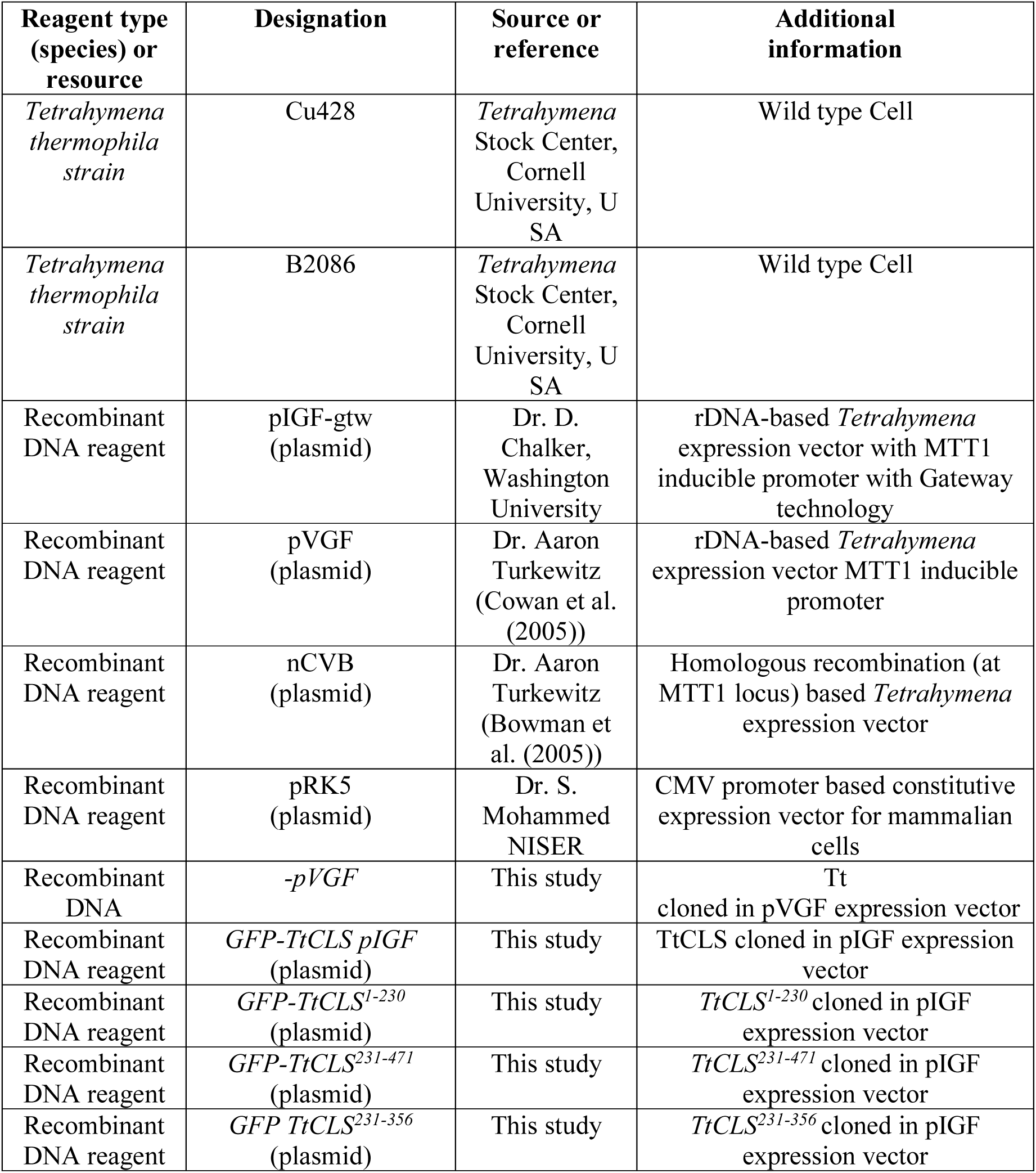
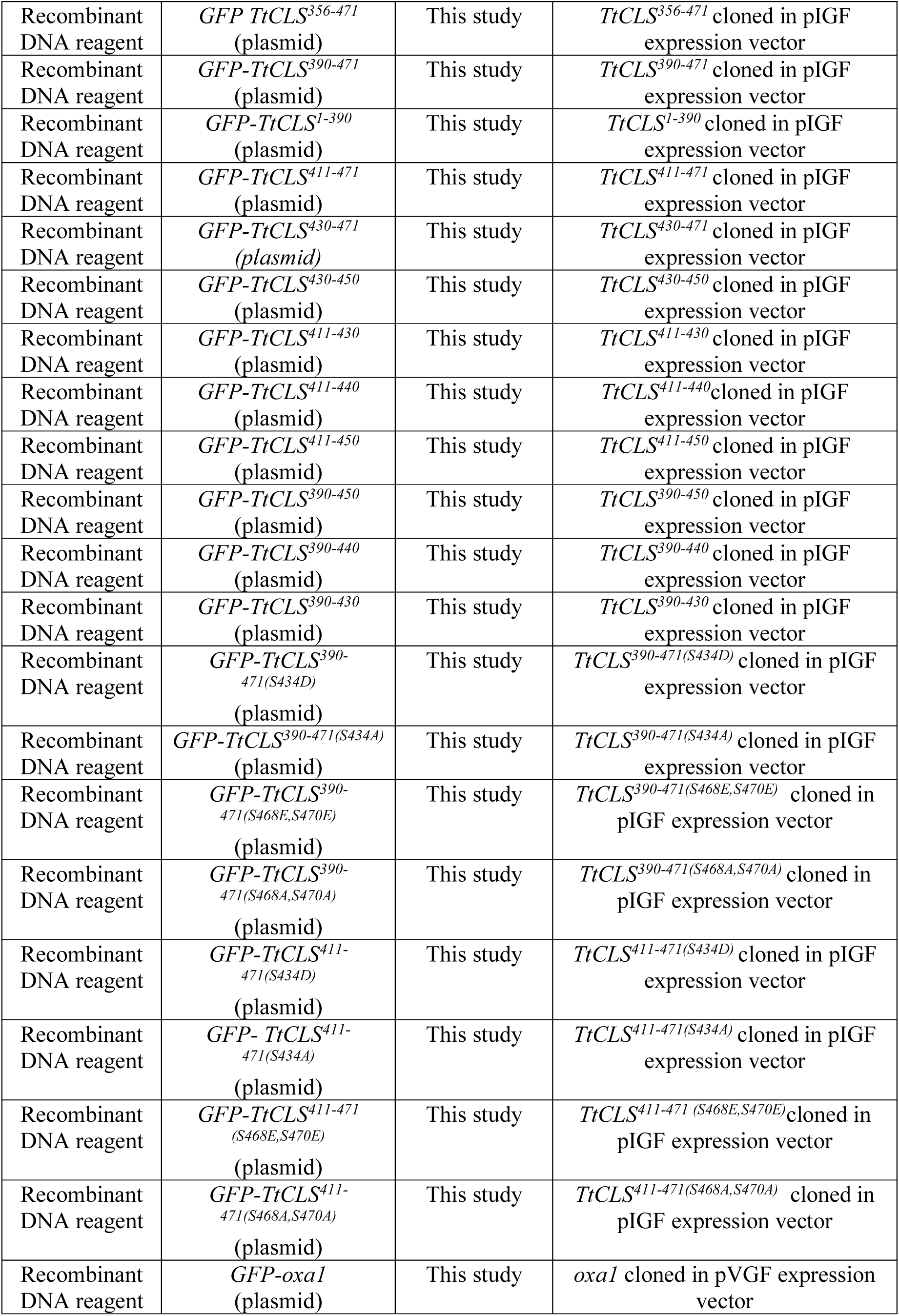
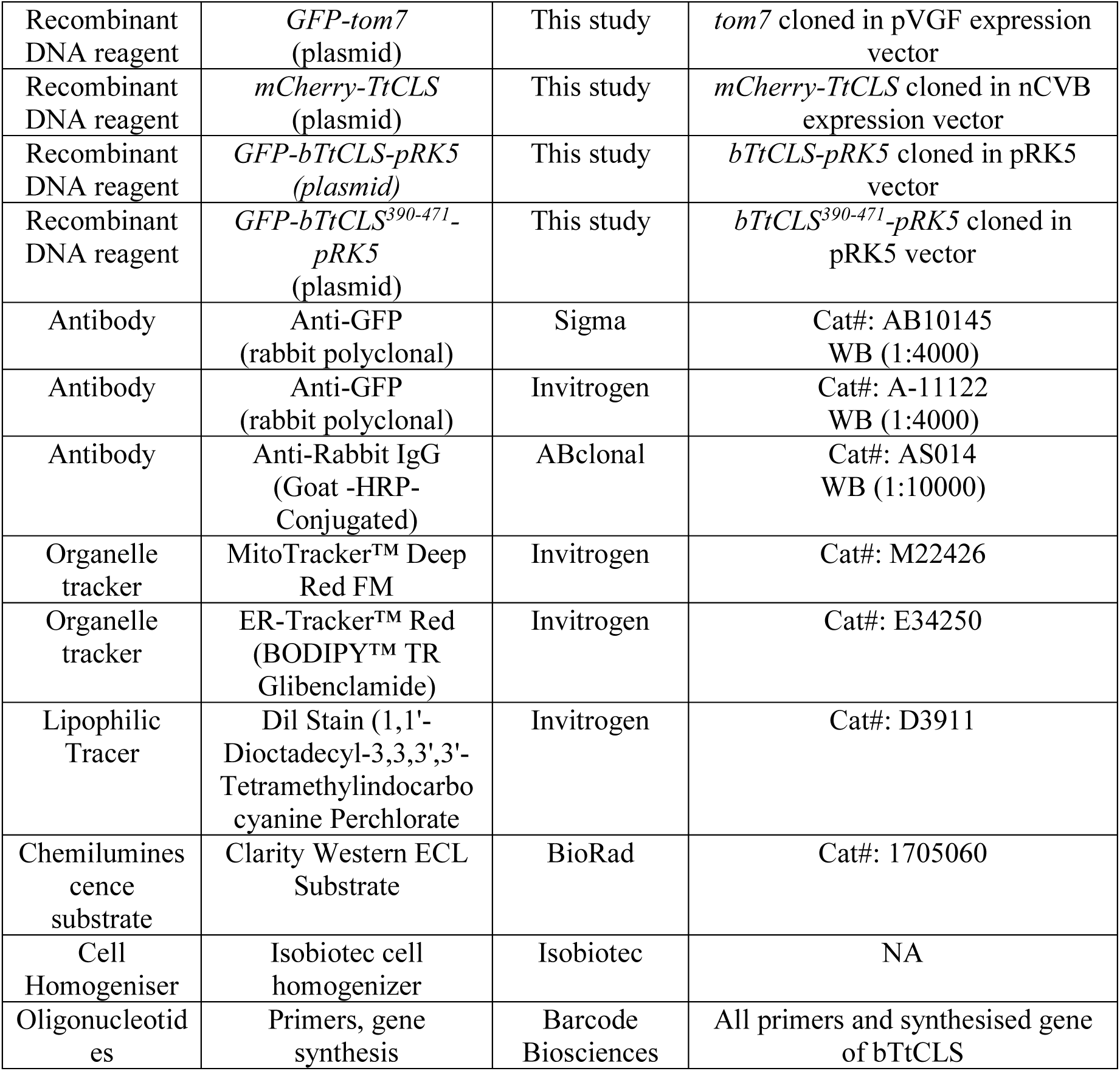

| **Reagent type (species) or resource** | **Designation** | **Source or reference** | **Additional information** |
|---|---|---|---|
| *Tetrahymena thermophila strain* | Cu428 | *Tetrahymena* Stock Center, Cornell University, USA | Wild type Cell |
| *Tetrahymena thermophila strain* | B2086 | *Tetrahymena* Stock Center, Cornell University, USA | Wild type Cell |
| Recombinant DNA reagent | pIGF-gtw (plasmid) | Dr. D. Chalker, Washington University | rDNA-based *Tetrahymena* expression vector with MTT1 inducible promoter with Gateway technology |
| Recombinant DNA reagent | pVGF (plasmid) | Dr. Aaron Turkewitz (Cowan et al. (2005)) | rDNA-based *Tetrahymena* expression vector MTT1 inducible promoter |
| Recombinant DNA reagent | nCVB (plasmid) | Dr. Aaron Turkewitz (Bowman et al. (2005)) | Homologous recombination (at MTT1 locus) based *Tetrahymena* expression vector |
| Recombinant DNA reagent | pRK5 (plasmid) | Dr. S. Mohammed NISER | CMV promoter based constitutive expression vector for mammalian cells |
| Recombinant DNA | *-pVGF* | This study | Tt cloned in pVGF expression vector |
| Recombinant DNA reagent | *GFP-TtCLS pIGF* (plasmid) | This study | TtCLS cloned in pIGF expression vector |
| Recombinant DNA reagent | *GFP-TtCLS<sup>1-230</sup>* (plasmid) | This study | *TtCLS<sup>1-230</sup>* cloned in pIGF expression vector |
| Recombinant DNA reagent | *GFP-TtCLS<sup>231-471</sup>* (plasmid) | This study | *TtCLS<sup>231-471</sup>* cloned in pIGF expression vector |
| Recombinant DNA reagent | *GFP TtCLS<sup>231-356</sup>* (plasmid) | This study | *TtCLS<sup>231-356</sup>* cloned in pIGF expression vector |
| Recombinant DNA reagent | *GFP TtCLS<sup>356-471</sup>* (plasmid) | This study | *TtCLS<sup>356-471</sup>* cloned in pIGF expression vector |
| Recombinant DNA reagent | *GFP-TtCLS<sup>390-471</sup>* (plasmid) | This study | *TtCLS<sup>390-471</sup>* cloned in pIGF expression vector |
| Recombinant DNA reagent | *GFP-TtCLS<sup>1-390</sup>* (plasmid) | This study | *TtCLS<sup>1-390</sup>* cloned in pIGF expression vector |
| Recombinant DNA reagent | *GFP-TtCLS<sup>411-471</sup>* (plasmid) | This study | *TtCLS<sup>411-471</sup>* cloned in pIGF expression vector |
| Recombinant DNA reagent | *GFP-TtCLS<sup>430-471</sup> (plasmid)* | This study | *TtCLS<sup>430-471</sup>* cloned in pIGF expression vector |
| Recombinant DNA reagent | *GFP-TtCLS<sup>430-450</sup>* (plasmid) | This study | *TtCLS<sup>430-450</sup>* cloned in pIGF expression vector |
| Recombinant DNA reagent | *GFP-TtCLS<sup>411-430</sup>* (plasmid) | This study | *TtCLS<sup>411-430</sup>* cloned in pIGF expression vector |
| Recombinant DNA reagent | *GFP-TtCLS<sup>411-440</sup>* (plasmid) | This study | *TtCLS<sup>411-440</sup>*cloned in pIGF expression vector |
| Recombinant DNA reagent | *GFP-TtCLS<sup>411-450</sup>* (plasmid) | This study | *TtCLS<sup>411-450</sup>* cloned in pIGF expression vector |
| Recombinant DNA reagent | *GFP-TtCLS<sup>390-450</sup>* (plasmid) | This study | *TtCLS<sup>390-450</sup>* cloned in pIGF expression vector |
| Recombinant DNA reagent | *GFP-TtCLS<sup>390-440</sup>* (plasmid) | This study | *TtCLS<sup>390-440</sup>* cloned in pIGF expression vector |
| Recombinant DNA reagent | *GFP-TtCLS<sup>390-430</sup>* (plasmid) | This study | *TtCLS<sup>390-430</sup>* cloned in pIGF expression vector |
| Recombinant DNA reagent | *GFP-TtCLS<sup>390-471(S434D)</sup>* (plasmid) | This study | *TtCLS<sup>390-471(S434D)</sup>* cloned in pIGF expression vector |
| Recombinant DNA reagent | *GFP-TtCLS<sup>390-471(S434A)</sup>* (plasmid) | This study | *TtCLS<sup>390-471(S434A)</sup>* cloned in pIGF expression vector |
| Recombinant DNA reagent | *GFP-TtCLS<sup>390-471(S468E,S470E)</sup>* (plasmid) | This study | *TtCLS<sup>390-471(S468E,S470E)</sup>* cloned in pIGF expression vector |
| Recombinant DNA reagent | *GFP-TtCLS<sup>390-471(S468A,S470A)</sup>* (plasmid) | This study | *TtCLS<sup>390-471(S468A,S470A)</sup>* cloned in pIGF expression vector |
| Recombinant DNA reagent | *GFP-TtCLS<sup>411-471(S434D)</sup>* (plasmid) | This study | *TtCLS<sup>411-471(S434D)</sup>* cloned in pIGF expression vector |
| Recombinant DNA reagent | *GFP- TtCLS<sup>411-471(S434A)</sup>* (plasmid) | This study | *TtCLS<sup>411-471(S434A)</sup>* cloned in pIGF expression vector |
| Recombinant DNA reagent | *GFP-TtCLS<sup>411-471 (S468E,S470E)</sup>* (plasmid) | This study | *TtCLS<sup>411-471 (S468E,S470E)</sup>*cloned in pIGF expression vector |
| Recombinant DNA reagent | *GFP-TtCLS<sup>411-471(S468A,S470A)</sup>* (plasmid) | This study | *TtCLS<sup>411-471(S468A,S470A)</sup>* cloned in pIGF expression vector |
| Recombinant DNA reagent | *GFP-oxa1* (plasmid) | This study | *oxa1* cloned in pVGF expression vector |
| Recombinant DNA reagent | *GFP-tom7* (plasmid) | This study | *tom7* cloned in pVGF expression vector |
| Recombinant DNA reagent | *mCherry-TtCLS* (plasmid) | This study | *mCherry-TtCLS* cloned in nCVB expression vector |
| Recombinant DNA reagent | *GFP-bTtCLS-pRK5 (plasmid)* | This study | *bTtCLS-pRK5* cloned in pRK5 vector |
| Recombinant DNA reagent | *GFP-bTtCLS<sup>390-471</sup>-pRK5* (plasmid) | This study | *bTtCLS<sup>390-471</sup>-pRK5* cloned in pRK5 vector |
| Antibody | Anti-GFP (rabbit polyclonal) | Sigma | Cat#: AB10145<br>WB (1:4000) |
| Antibody | Anti-GFP (rabbit polyclonal) | Invitrogen | Cat#: A-11122<br>WB (1:4000) |
| Antibody | Anti-Rabbit IgG (Goat -HRP-Conjugated) | ABclonal | Cat#: AS014<br>WB (1:10000) |
| Organelle tracker | MitoTracker™ Deep Red FM | Invitrogen | Cat#: M22426 |
| Organelle tracker | ER-Tracker™ Red (BODIPY™ TR Glibenclamide) | Invitrogen | Cat#: E34250 |
| Lipophilic Tracer | Dil Stain (1,1'-Dioctadecyl-3,3,3',3'-Tetramethylindocarbocyanine Perchlorate | Invitrogen | Cat#: D3911 |
| Chemiluminescence substrate | Clarity Western ECL Substrate | BioRad | Cat#: 1705060 |
| Cell Homogeniser | Isobiotec cell homogenizer | Isobiotec | NA |
| Oligonucleotides | Primers, gene synthesis | Barcode Biosciences | All primers and synthesised gene of bTtCLS |

## List of primers

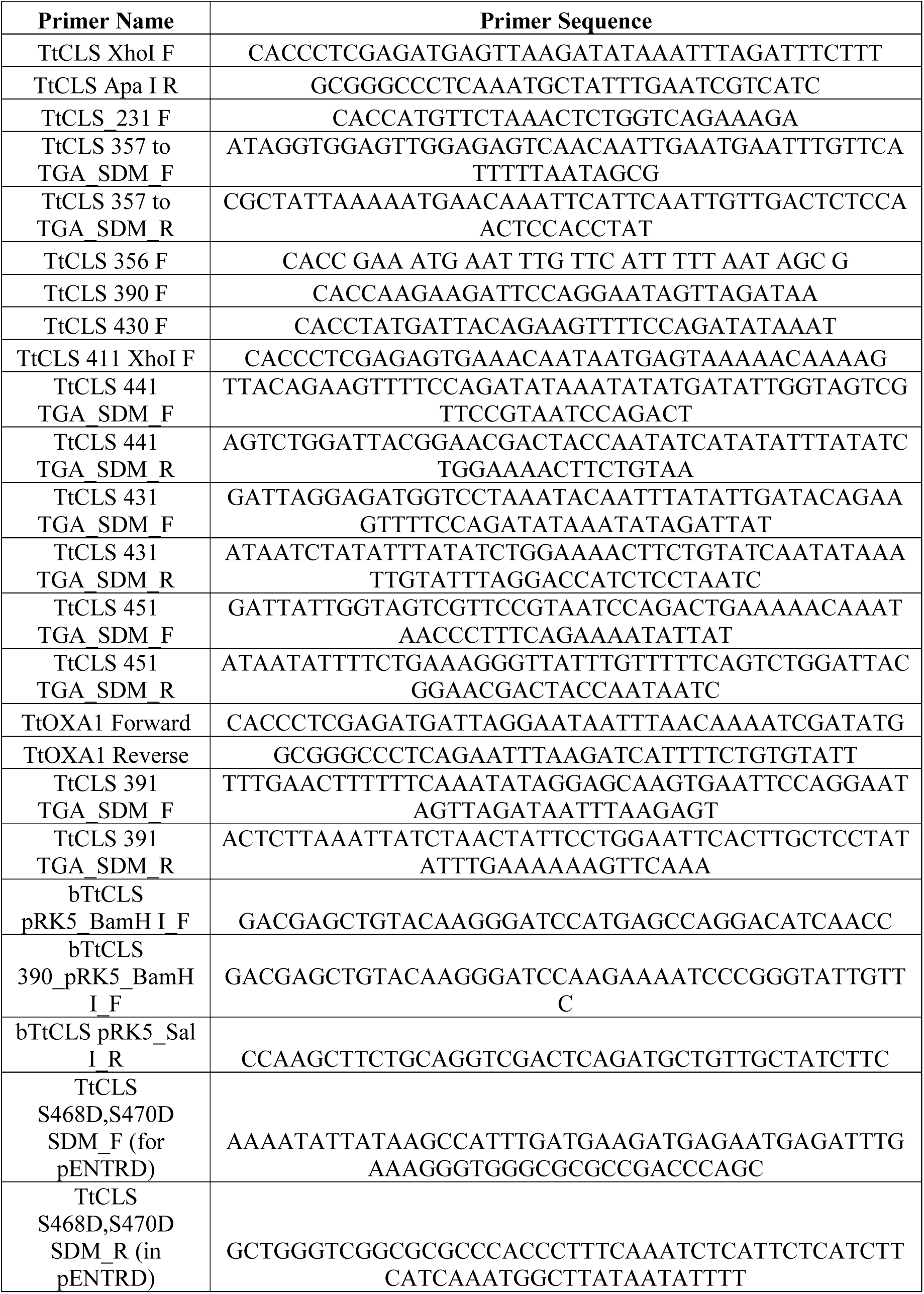
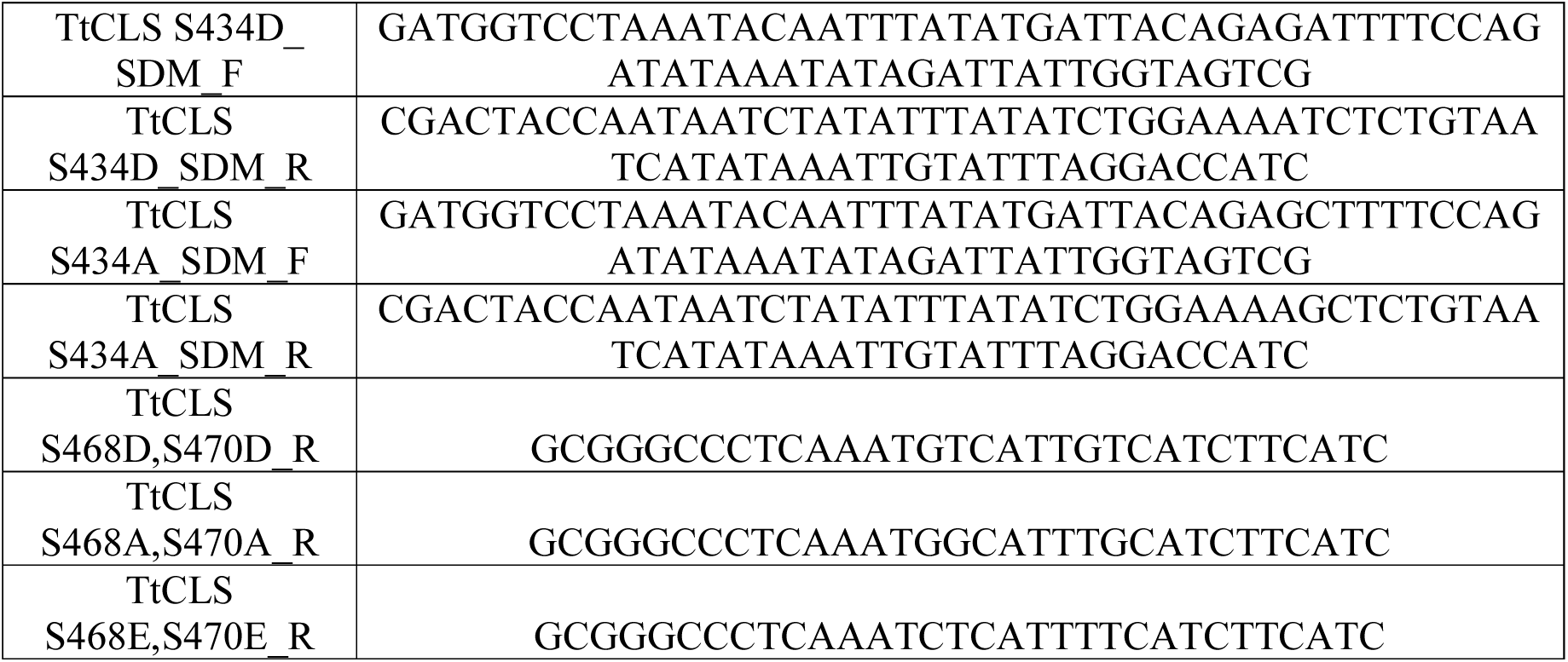

